# *Xrp1* drives damage-induced cellular plasticity of enteroendocrine cells in the adult *Drosophila* midgut

**DOI:** 10.1101/2025.07.05.662934

**Authors:** Qingyin Qian, Hiroki Nagai, Yuya Sanaki, Makoto Hayashi, Kenichi Kimura, Yu-ichiro Nakajima, Ryusuke Niwa

**Author notes:** Corresponding authors’ email addresses (Q.Q.) (R.N.). Institute for Aquaculture Biotechnology, Tokyo University of Marine Science and Technology, 4-5-7 Konan, Minato-ku, Tokyo, 108-8477, Japan (Makoto Hayashi).

## Abstract

Cellular plasticity, the ability of a differentiated cell to adopt another phenotypic identity, is restricted under basal conditions, but can be elicited upon damage to facilitate regeneration. Such damage-induced cellular plasticity restores homeostasis and prevents pathology, yet its underlying molecular basis remains largely unexplored. Here, we reported damage-induced cellular plasticity of secretory enteroendocrine cells (EEs) in the adult *Drosophila* midgut. We found that ionizing radiation enhanced EE plasticity such that it promoted EEs to dedifferentiate into ISCs and subsequently re-differentiate towards ECs. We identified that radiation induced the production of a stress-inducible transcription factor Xrp1 in EE lineages, and its upregulation was necessary for EE plasticity. Single-cell RNA sequencing of guts with EE-specific *Xrp1* overexpression revealed ectopic expression of progenitor-specific genes in EEs, which was necessary for *Xrp1* to drive EE plasticity. Our work provides a mechanistic framework for understanding cellular plasticity and suggests its potential role in damage-induced responses.

## Introduction

Cellular plasticity is defined as the ability of a differentiated cell to change its phenotypic identity. As a cell progresses through differentiation, its identity becomes solidified to ensure optimal functionality, and its plasticity becomes constrained to preserve tissue architecture. While cell identity is restricted under normal conditions, differentiated cells are not always locked into their fate but exhibit a certain degree of plasticity, enabling them to undergo dedifferentiation and transdifferentiation. A well-studied *in vitro* example is the generation of induced pluripotent stem cells from fully differentiated cells (Takahashi et al., 2007; J. Yu et al., 2007). Cellular plasticity is further reinforced by multiple lines of *in vivo* evidence, which tie cellular plasticity to damage-induced tissue repair. For instance, differentiated epithelial cells can revert to a stem-like state to regenerate the damaged airway and intestine (Adkins-Threats & Mills, 2022; Kim et al., 2024; Murata et al., 2020; Tata et al., 2013). Hepatocytes and biliary epithelial cells can convert fate to one another upon liver injury, which contributes to the liver’s remarkable regenerative capacity (Li et al., 2020; Schaub et al., 2018). Notably, aberrant cellular plasticity has been linked to disease development, particularly cancer progression (Pérez-González et al., 2023). These observations underscore the necessity of elucidating the underlying mechanism and the physiological significance of cellular plasticity, especially in the context of damage-induced regeneration.

The intestinal epithelial cell lineage serves as an ideal *in vivo* model for studying such cellular plasticity. In mammals, intestinal stem cells (ISCs) differentiate to produce both absorptive and secretory cell lineages, while upon perturbation, cells from both lineages can revert backwards to replenish ISCs (Jones et al., 2019; Tetteh et al., 2016; H. Tian et al., 2011; van Es et al., 2012; Yan et al., 2017; S. Yu et al., 2018). The cell type composition and lineage of intestinal epithelial cells are highly conserved yet much simpler in the model organism, fruit fly *Drosophila melanogaster* (Miguel-Aliaga et al., 2018). Within the adult *Drosophila* intestinal epithelial cell lineage, ISCs undergo asymmetric division to produce a renewed ISC and a committed progenitor cell, either an enteroblast (EB) or an enteroendocrine progenitor cell (EEP), which thereafter respectively gives rise to an absorptive enterocyte (EC) with a large polyploid nucleus, or divides once to yield a pair of secretory enteroendocrine cells (EEs) with a small diploid nucleus (Chen et al., 2018; Micchelli & Perrimon, 2006; Ohlstein & Spradling, 2006; Zeng & Hou, 2015). Importantly, recent studies have provided key insights into the cellular plasticity of *Drosophila* intestinal epithelial cells. For example, forced depletion of *tramtrack* (*ttk*) can trans-differentiate ECs into Ees, while forced depletion of *prospero* (*pros*) can dedifferentiate EEs into ISCs (Guo et al., 2024). Bacterial infection triggers EBs to enter mitosis and produce ISCs to replenish the ISC pool (A. Tian et al., 2022), while fluctuations in nutrient availability induce EEs to dedifferentiate into ISCs, which then more preferentially re-differentiate into ECs to adaptively resize the shrunken midgut (Nagai et al., 2023). However, a comprehensive analysis of EE plasticity, both under basal conditions and in the context of damage-induced responses, has not yet been conducted.

In this study, we started from the observation of baseline EE plasticity being enhanced by radiation and uncovered a stress-activated transcription factor *Xrp1* (Cong & Cagan, 2024), which was inducible in EE lineages by radiation and necessary for radiation-induced EE plasticity. Furthermore, our findings added a new role of *Xrp1* in regulating the expression of progenitor-specific genes, whose ectopic expression in EEs was necessary for *Xrp1*-driven EE plasticity. By uncovering the molecular framework governing radiation-induced EE plasticity, we provide broader insights into epithelial damage responses and propose the possibility that such plasticity contributes to tissue repair.

## Results

### EEs exhibit cellular plasticity under baseline conditions

To examine EE plasticity in the adult *Drosophila* midgut, we performed lineage tracing of EEs using an EE-specific Gal4 driver, *prospero-Gal4* (*pros-Gal4*), combined with a temperature-sensitive form of the Gal80 repressor (*tub-Gal80^ts^*), to the lineage trace tool G-TRACE (Evans et al., 2009), which allowed us to distinguish the current expression (RFP) and the previous expression (GFP) of the Gal4 driver. With this system, current *pros*-expressing EEs are marked with GFP and RFP, whereas cells that were EEs and expressed *pros* are marked with GFP solely and permanently (Fig. 1A). We first confirmed that this system exhibited minor leaky labeling under restrictive temperatures (Fig. S1A). We initiated EE lineage tracing within 2 days of adult eclosion and collected midguts from these flies after 7 days. Immunostaining of these guts reassured us that RFP^+^ GFP^+^ cells and RFP^+^ GFP^weak^ cells, which likely had not yet fully activated recombination, indeed colocalized with the nucleus-localized EE marker Prospero (Pros); hence, we referred to them as current EEs. On the contrary, RFP^weak^ GFP^+^ cells and RFP^-^ GFP^+^ cells surely lost the EE marker Pros but rather expressed the puncta-shaped cytoplasm-localized ISC marker Delta (Dl); hence, we referred to them as past EEs (Fig. 1B). Additionally, RFP^-^ GFP^weak^ small cells were observed, whose GFP intensity was much weaker than that of current EEs and past EEs. These cells were negative for the EE marker Pros but positive for the ISC marker Dl (Fig. 1B). We assumed these cells were ISCs that had expressed *pros* transiently, whose level was too weak to be committed into EEPs (Chen et al., 2018; Wu et al., 2023); hence, we did not include these cells in EE lineages.

**Figure 1.**
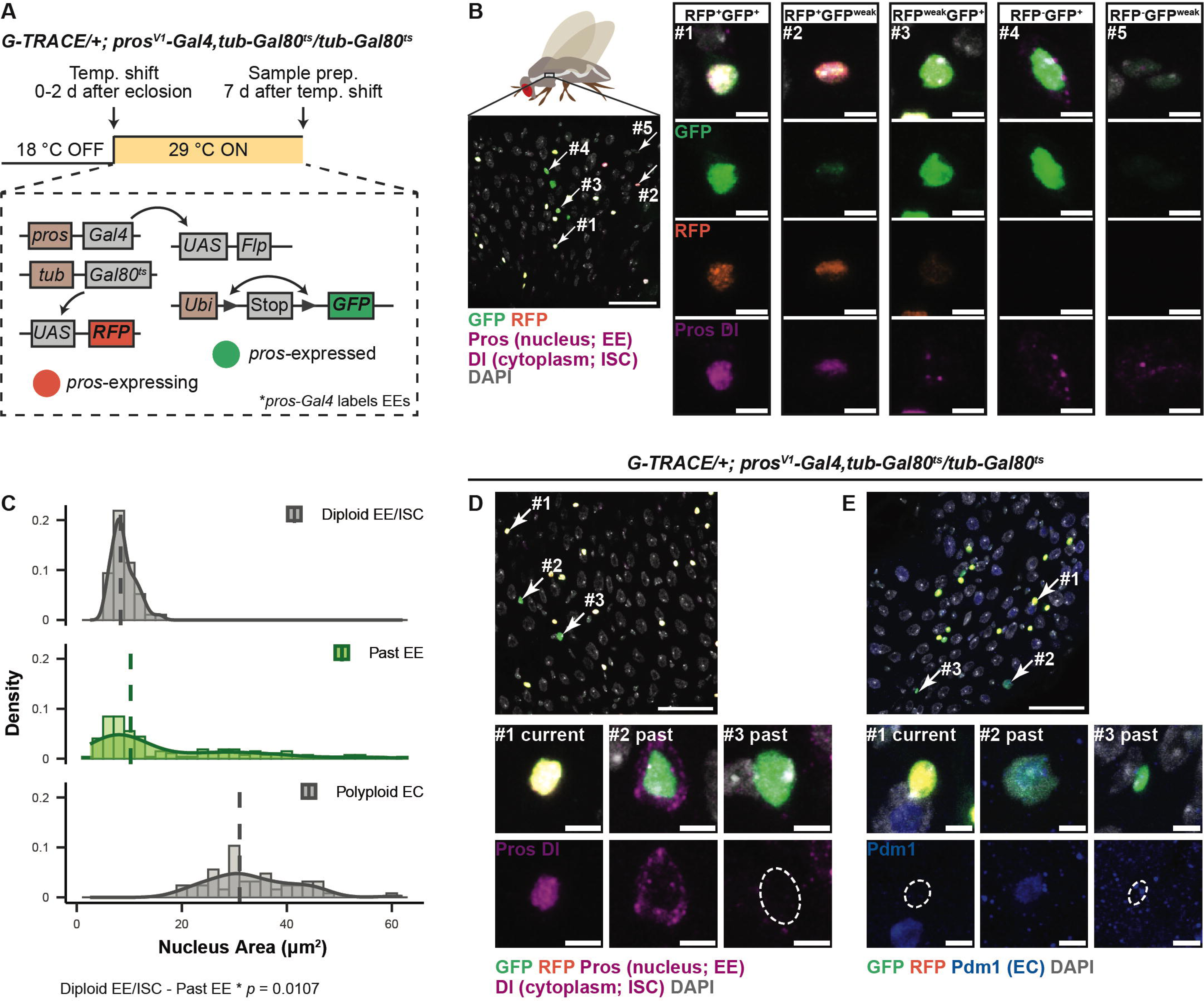
**EEs exhibit cellular plasticity and can alter their identity into ISCs or ECs under baseline conditions.** (A) Schematic of EE lineage tracing. Given that *pros-Gal4* specifically targets EEs, crossing *pros-Gal4* to G-TRACE marks *pros*-expressing EEs with RFP, and cells had *pros* expressed with GFP permanently. (B) Left: Representative EE lineage-traced gut image; stained with anti-GFP (green), anti-RFP (orange), anti-Pros (magenta; nucleus-localized), anti-Dl (magenta; cytoplasm-localized), and DAPI (white). Scale bar: 30 μm. Right: Magnified views of cells indicated by numbered arrows on the left image. Top: merged; Second: GFP; Third: RFP; Bottom: Pros and Dl. Scale bars: 3 μm. (C) Quantification of the nucleus area of diploid EEs/ISCs, past EEs, and polyploid ECs; illustrated by density plots showing the distribution and dashed lines indicating medians. N[gut] = 32, n[past EE] = 136, n[EE/ISC] = 48, n[EC] = 64. Kruskal-Wallis test followed by Steel-Dwass Test. (D and E) Top: Representative EE lineage-traced gut images; both stained with anti-GFP (green), anti-RFP (orange), and DAPI (white); additionally, (D) stained with anti-Pros (magenta; nucleus-localized) and anti-Dl (magenta; cytoplasm-localized), (E) stained with anti-Pdm1 (blue). Scale bars: 30 μm. Bottom: Magnified views of cells indicated by numbered arrows on the image above. Top: merged; Bottom: (D) Pros and Dl; (E) Pdm1. Scale bars: 3 μm.

Across the regionally defined adult *Drosophila* midgut (Buchon et al., 2013), past EEs in the anterior (R2a, R2b, and R2c) and middle (R3) midgut all appeared to be dispersed rather than clustered, while those in the posterior (R4a and R4b) midgut more seemingly appeared clustered, indicative of a group of cells descended from an EE-derived ISC (Fig. S1B). As the relatively low cell density allowed more accurate quantification, we thereafter focused on the anterior midgut and calculated the percentage of past EEs among *pros*-expressing and -expressed EE lineages per region of interest (ROI). We found that past EEs exhibited differential representation between R2a and R2b. In the R2a region, past EEs constituted 5.18 ± 0.89 % of EE lineages, surpassing what was found in the R2b region, where they only comprised 1.78 ± 0.36 % of EE lineages (Fig. S1C). Although the exact reason for this occurrence remained uncertain, we limited our future analysis to the R2a region. Because *pros-Gal4* labels EE progenitor (EEPs) (Guo et al., 2019), in addition to mature EEs, we used another EE-specific driver, *Neuropeptide F-T2A-Gal4* (*NPF-T2A-Gal4*), which more specifically targets differentiated EEs, to confirm the reproducibility of EE fate conversion (Fig. S1D). Overall, these results highlight that a small subset of post-mitotic EEs are plastic, such that they can adopt another cell fate.

### Under basal conditions, EEs primarily convert into ISCs, which differ from normal ISCs

In the adult *Drosophila* midgut epithelium, cells can be distinguished by their nuclear size and molecular identity. ISCs, EBs, and EEs are small and diploid, whereas ECs undergo endoreplication to become large and polyploid (Øvrebø & Edgar, 2018). To accurately address the identity of EE-derived past EEs, we first measured their nuclear size. Past EEs exhibited a primary peak of nuclear size characteristic of diploid EEs/ISCs, but a subset showed a lagging distribution with nuclear sizes resembling those of polyploid ECs, implying a heterogeneous composition of ISCs, EBs, and ECs (Fig. 1C). We thus examined the possibility of past EEs colocalizing with cell type markers. While current EEs overlapped with the nucleus-localized EE marker Pros without cytoplasm-localized ISC marker Dl, 55.3 % (26/47) past EEs rather colocalized with the ISC marker Dl (Fig. 1D). On the other hand, 36.5 % (27/74) past EEs colocalized with the EC marker Pdm1 (Fig. 1E). We next examined another progenitor feature, a high-level and pointed-shaped adherens junction, visualized by Armadillo (Arm) (Singh et al., 2012). Some past EEs of a small nuclear size featured high levels of sharp triangular Arm signals characteristic of progenitor cells, while others retained EE-like rounded Arm signals, indicating an EE-to-ISC transition (Fig. S1E) (Nagai et al., 2023). Past EEs of a large nuclear size and their adjacent ECs had septate junction complex protein, indicated by Coracle (Cora) (Fig. S1F), suggesting proper integration of such past EEs into the gut epithelium (Furuse & Izumi, 2017). These observations reinforced the idea that past EEs correctly adopt distinct cell identity, primarily ISCs, with the potential to become ECs.

To precisely examine the molecular signature of past EEs and possibly distinguish them from normally differentiated cells, we combined the lineage tracing with single-cell RNA sequencing (scRNA-seq). We generated two datasets of EE lineages (both RFP^+^ GFP^+^ current EEs and RFP^-^ GFP^+^ past EEs) by sorting GFP^+^ cells via Fluorescence-Activated Cell Sorting (FACS), and one dataset from unsorted EE lineage-traced guts, encompassing all cell types. The three datasets were filtered, integrated, and projected onto the uniform manifold approximation and projection (UMAP) plot. Cell types were determined by established marker gene expression (Guo et al., 2019; Hung et al., 2020), and each EE cluster was further refined by annotating it according to the major peptide hormone, such as Allatostatins (AstA, AstB/Mip, AstC), Tachykinin (Tk), or CCHamide-1 (CCHa1), that showed the highest expression in that cluster (Fig. S1G). Subsequently, we focused on the _“_ISC/EB_”_, defined by the progenitor marker *escargot* (*esg*) expression. Within this population, a subset was *RFP*^-^ *GFP*^+^, lacking the EE marker *pros* expression, indicative of past EEs, thereby being relevant to our primary objective (Fig. S1H). Marker gene analysis between *GFP*^+^ EE-derived _“_ ISC/EB_”_ and *GFP*^-^ normal _“_ ISC/EB_”_ revealed that a transcription factor, *ets21C*, implicated in stress-induced regenerative responses by promoting ISC proliferation and differentiation (Mundorf et al., 2019), was significantly enriched in EE-derived _“_ISC/EB_”_, despite being under normal conditions (Fig. S1I). This difference suggests that in homeostatic guts, dedifferentiation can be triggered by stress or accompanied by stress-related genetic programs, and EE-derived progenitor cells are in a regenerative state with increased proliferative capacity and differentiation potential.

### Ionizing radiation (IR) exposure induces EEs to gain plasticity towards EC fate

It is known that ISC proliferation and differentiation are the primary mechanisms by which the *Drosophila* gut epithelium repairs itself following damage (Amcheslavsky et al., 2009; Biteau et al., 2008; Buchon et al., 2009; Jiang et al., 2011). Besides ISC activities, we questioned whether EE plasticity was also involved in damage responses. First, we tested commonly used gut-damaging chemicals, Dextran Sulfate Sodium (DSS) and paraquat. However, these treatments did not alter the number of past EEs (Fig. S2A – S2B). We next explored the effect of ionizing radiation (IR), which generates DNA damage and disrupts homeostatic intestinal epithelium (Huang & Zhou, 2020; Stacey & Green, 2014). We subjected flies to 100 Gy of X-ray irradiation 6 days after initiation of lineage tracing and collected their midguts several hours after radiation on day 6 or day 7 (Fig. 2A). A time-course analysis highlighted that the number of past EEs increased dramatically, peaked 10-14 hours after radiation and decreased thereafter, while the number of current EEs remained unchanged by radiation (Fig. 2B – 2C). These results indicate that radiation triggers EE plasticity, and this transformation of EE identity is a transient event. In addition, consistent with the previous study (Sharma et al., 2020), we observed a decrease in the number of ISCs in irradiated guts; notably, we confirmed that it occurred within hours after radiation (Fig. S2C). The timing of the ISC decrease coincided with the induction of EE plasticity, implying a potential correlation between the two events.

**Figure 2.**
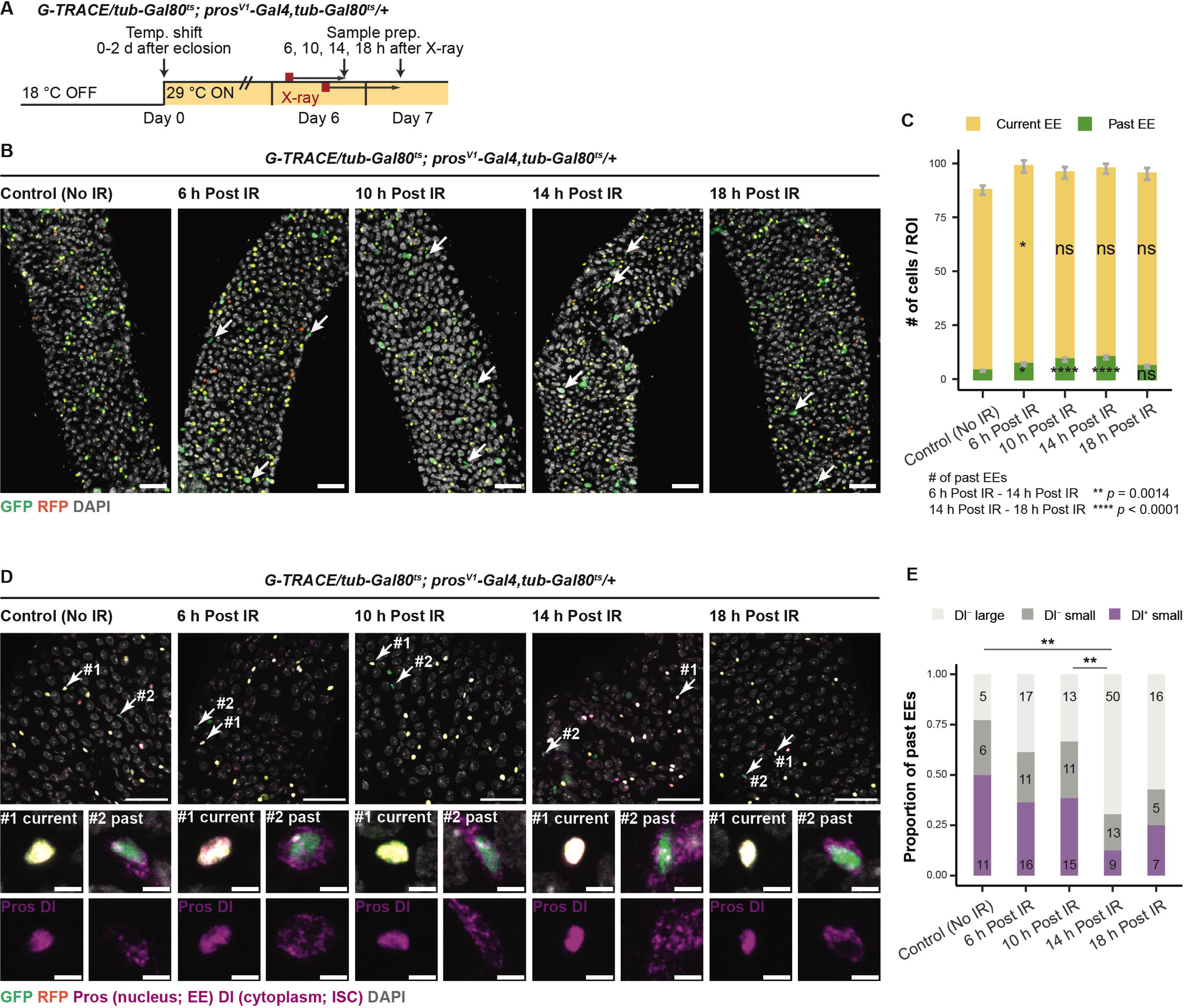
**Ionizing irradiation (IR) promotes EEs to gain plasticity towards EC fate.** (A) Schematic of radiation exposure to EE lineage-traced flies. (B) Representative EE lineage-traced gut images of non-irradiated (No IR) and irradiated (6 h, 10 h, 14 h, and 18 h Post IR) flies (from left to right); stained with anti-GFP (green), anti-RFP (orange), and DAPI (white); past EEs highlighted by red arrows. Scale bars: 30 μm. (C) Quantification of the number of past EEs and current EEs in non-irradiated (No IR) and irradiated (6 h, 10 h, 14 h, and 18 h Post IR) guts, corresponding to (B). N[gut] = 37 (No IR), 39 (6 h Post IR), 38 (10 h Post IR), 48 (14 h Post IR), 38 (18 h Post IR). Current EE: one-way ANOVA followed by Tukey HSD, * *p* < 0.05; past EE: Kruskal-Wallis test followed by Steel-Dwass Test, * *p* < 0.05, **** *p* < 0.0001. (D) Top: Representative EE lineage-traced gut images of non-irradiated (No IR) and irradiated (6 h, 10 h, 14 h, and 18 h Post IR) flies (from left to right); stained with anti-GFP (green), anti-RFP (orange), anti-Pros (magenta; nucleus-localized), anti-Dl (magenta; cytoplasm-localized), and DAPI (white). Scale bars: 30 μm. Bottom: Magnified views of cells indicated by numbered arrows on the image above. Top: merged; Bottom: Pros and Dl. Scale bars: 3 μm. (E) The proportion of past EEs being small or large in nucleus size and being positive or negative for the ISC marker Dl, between non-irradiated (No IR) and irradiated (6 h, 10 h, 14 h, and 18 h Post IR) guts, corresponding to (D). Pairwise Fisher’s exact test, ** *p* < 0.01.

In addition to examining radiation-induced changes in EE plasticity, as indicated by the number of current and past EEs, we also investigated whether there was any change in the cell type composition of past EEs. Co-staining of the ISC marker Dl showed that the likelihood of past EEs being Dl^+^ ISCs remained unchanged between non-irradiated control guts and irradiated guts examined 6 or 10 hours after radiation, while in the guts examined 14 hours after radiation, past EEs had a smaller chance of being Dl^+^ ISCs (Fig. 2D – 2E). Supporting this, nuclear size measurements revealed that past EEs in the guts examined 14 hours after radiation, as compared to those in non-irradiated guts, shifted towards having a larger nucleus, which was indicative of EC identity (Fig. S2D). Together, these results indicate that EEs exhibit enhanced plasticity in response to radiation, with a greater number of EEs dedifferentiating into ISCs and subsequently re-differentiating towards an EC fate.

### *Xrp1* upregulation accounts for radiation-induced EE plasticity

Having observed radiation-induced EE plasticity, we questioned whether there could be any transcriptional changes driving this process. It has been known that *Xrp1*, a stress-inducible transcription factor, is upregulated in *Drosophila* embryos and wing discs after X-ray irradiation (Akdemir et al., 2007; Brodsky et al., 2004; Cruz et al., 2025). We thus examined whether *Xrp1* could be induced in irradiated guts. In wild-type flies, anti-Xrp1 staining was uniformly absent in non-irradiated guts but present in some cells in irradiated guts. Notably, this signal was not seen in irradiated *Xrp1* knock-out flies, confirming antibody specificity (Fig. S3A) (Mallik et al., 2018). We then returned to EE lineage-traced flies to address whether EE lineages were where *Xrp1* induction occurred. We observed that Xrp1 protein was produced in 23.5 ± 4.0% of current EEs and 40.96% (34/83) of past EEs after irradiation (Fig. 3A–3C). Meanwhile, Xrp1 protein was detected in part of non-EE small cells that were expected to be progenitor cells, and part of large cells that were expected to be ECs in irradiated guts (Fig. S3B). Using *Xrp1-LacZ* that reflects *Xrp1* transcripts, we confirmed a radiation-induced increase in the ratio of cells that were positive for *Xrp1-LacZ* in both Pros^+^ EEs and Dl^+^ ISCs (Fig. S3C – S3E). These data suggest that *Xrp1* mRNA and Xrp1 protein can be upregulated in all cell types, including current and past EEs, following radiation.

**Figure 3.**
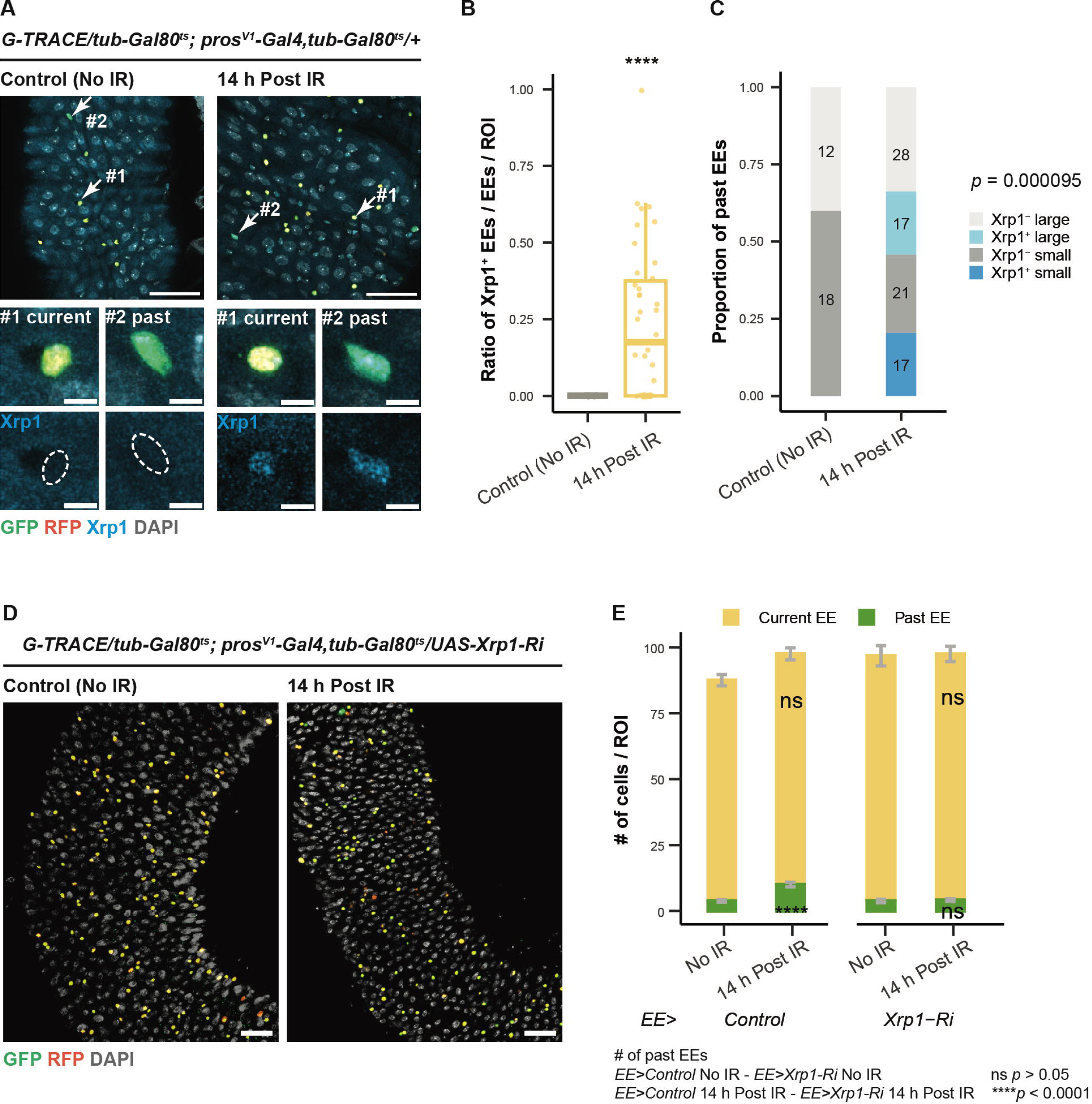
***Xrp1* upregulation in EEs is essential for radiation-induced EE plasticity.** (A) Top: Representative EE lineage-traced gut images of non-irradiated (No IR; top left) and irradiated (14 h Post IR; top right) flies; stained with anti-GFP (green), anti-RFP (orange), anti-Xrp1 (cyan), and DAPI (white). Scale bars: 30 μm. Bottom: Magnified views of cells indicated by numbered arrows on the image above. Top: merged; Bottom: Xrp1. Scale bars: 3 μm. (B) Quantification of the ratio of Xrp1^+^ EEs among EEs per ROI, corresponding to (A). N[image] = 17 (No IR), 38 (14 h Post IR). Wilcoxon Rank-Sum test, **** *p* < 0.0001. (C) The proportion of past EEs being small or large in nucleus size and being positive or negative for Xrp1, between non-irradiated and irradiated (14 h Post IR) guts. Fisher’s exact test. (D) Representative EE lineage-traced gut images of non-irradiated (No IR; left) and irradiated (14 h Post IR; right) flies with *Xrp1* knocked down in EEs; stained with anti-GFP (green), anti-RFP (orange), and DAPI (white). Scale bars: 30 μm. (E) Quantification of the number of past EEs and current EEs, corresponding to (D). *EE > Control*: N[gut] = 37 (No IR), 48 (14 h Post IR). *EE >Xrp1-Ri*: N[gut] = 20 (No IR), 30 (14 h Post IR). Current EE: one-way ANOVA followed by Tukey HSD; past EE: Kruskal-Wallis test followed by Steel-Dwass Test, **** *p* < 0.0001. *EE > Control* No IR and 14 h Post IR data were reproduced from Fig. 2C to aid comparison.

Considering the induction of *Xrp1* mRNA and Xrp1 protein in EEs and their fate conversion after radiation, we then questioned if *Xrp1* was required for radiation-induced EE plasticity by depleting *Xrp1* in EEs. We first assessed *Xrp1* knockdown efficiency by checking Xrp1 protein production and confirmed the decrease in the ratio of EEs positive for Xrp1 protein in irradiated *Xrp1* knocked-down guts (Fig. S3E). While *Xrp1* knockdown in EEs under normal conditions did not affect the number of current and past EEs, knocking down *Xrp1* in EEs prevented the increase of past EEs in irradiated guts (Fig. 3D – 3E), indicating the essential role of *Xrp1* in radiation-induced EE plasticity. While *Xrp1* is required for apoptosis of loser cells in cell competition (Baillon et al., 2018), considering that the EE number was not reduced shortly after radiation (Fig. 3E), we could rule out the possibility that apoptosis prevention accounted for the inhibition of radiation-induced EE plasticity upon *Xrp1* knockdown. In contrast, 7 days after radiation, the EE number was reduced by irradiation, and this reduction was blocked upon *Xrp1* knockdown in EEs (Fig. S3F – S3G), implying Xrp1’s established apoptotic functions. Altogether, we conclude that radiation-responsive *Xrp1* upregulation in EE lineages is essential for radiation-induced EE plasticity.

### Forced *Xrp1* expression in EEs imposes progenitor identities on EEs

So far, we identified *Xrp1* upregulation in EEs as a critical driver in radiation-induced EE plasticity. To further examine transcriptional changes resulting from *Xrp1* upregulation in EEs, we overexpressed *Xrp1* in EEs under basal conditions (*Xrp1* O/E EE) and performed scRNA-seq of the anterior and middle midguts under control and *Xrp1* O/E EE conditions. With marker gene-based approaches, we identified one ISC/EB cluster, four anterior-midgut EC (aEC) clusters, one middle-midgut EC (mEC) cluster and three EE clusters and excluded cardia cell clusters, as shown in the UMAP plot (Fig. 4A). To get a clearer view of EE subtypes, EE 1-3 were clusters subset and re-categorized, resulting in five EE clusters (Fig. 4B – 4C): _“_ AstC.EE_”_ , _“_ CCHa2.EE_”_ and _“_ Tk.EE_”_ named after their relatively highly expressed peptide hormones, _“_esg.EE_”_ named due to *esg* expression, which was expected to represent middle-midgut EEs (Hung et al., 2020), _“_N.EE_”_ named due to progenitor marker *Notch* (*N*) expression, which was expected to represent cells in the transition of either ISC-EE differentiation (Guo et al., 2019) or EE-ISC dedifferentiation (Nagai et al., 2023).

**Figure 4.**
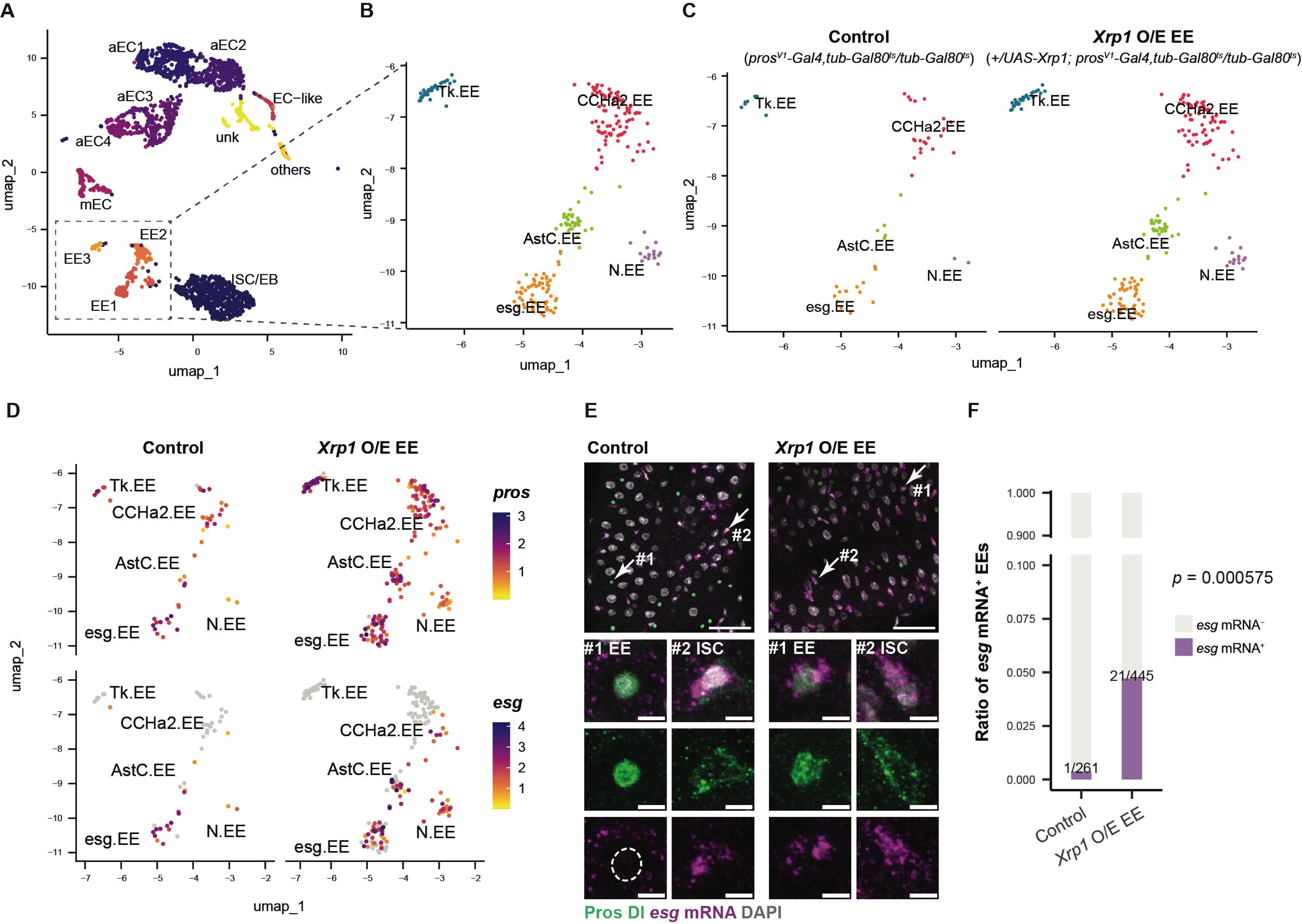
***Xrp1* O/E induces progenitor cell-specific gene expression in EEs.** (A) UMAP plot of control guts and guts with *Xrp1* O/E in EEs. (B) UMAP plot of re-clustered EE clusters. (C) UMAP plot of re-clustered EE clusters split by the genotype. (D) Feature plot showing normalized expression of *pros* and *esg* in each EE cluster split by the genotype (left: control, right: *Xrp1* O/E EE; top: *pros*, bottom: *esg*). (E) Top: Representative gut images of control (top left) and *Xrp1* O/E in EEs (top right); stained with anti-Pros (green; nucleus-localized), anti-Dl (green; cytoplasm-localized), *esg* mRNA (magenta; cytoplasm-localized), and DAPI (white). Scale bars: 30 μm. Bottom: Magnified views of cells indicated by numbered arrows on the image above. Top: merged; Middle: Pros and Dl; Bottom: *esg* mRNA. Scale bars: 3 μm. (F) Quantification of the ratio of *esg* mRNA^+^ Pros^+^ EEs among Pros^+^ EEs. Fisher’s exact test.

To identify gene expression changes driven by *Xrp1*, we investigated how *Xrp1* O/E in EEs altered EEs’ gene expression profiles. We first checked EEs’ peptide hormone expression levels to see if there were any differences in EE identity. We noticed a decreased *AstC* expression in _“_AstC.EE_”_, a decreased *Tk* expression in _“_Tk.EE_”_, and a decreased *NPF* expression in _“_N.EE_”_, suggesting that *Xrp1* O/E in EEs diminishes their EE identity (Fig. S4A). We then assessed changes in cell type markers. As a result, progenitor cell-specific transcription factors that are known to maintain ISC identity or prime ISCs towards ECs, *esg*, *Sox100B*, *Sox21a* and *klumpfuss* (*klu*), were ectopically expressed in part of _“_AstC.EE_”_, _“_CCHa2.EE_”_, and _“_Tk.EE_”_ (Fig. 4D, S4B, and S4C) (Antonello et al., 2015; Chen et al., 2016; Jin et al., 2020; Korzelius et al., 2019, 2014), suggesting that *Xrp1* O/E in EEs forces EEs to gain ISC identity, which is in turn destined to acquire EC identity. To validate this notion, we visualized *esg* mRNA and examined its colocalization with the EE marker Pros. In the anterior midgut, *esg* mRNA was consistently observed in Dl^+^ ISCs but rarely observed in Pros^+^ EEs (0.38 %, 1/261) in control guts; in contrast, in *Xrp1* O/E EE guts, an increased incidence of Pros^+^ EEs expressing *esg* mRNA (4.7 %, 21/445) was noted (Fig. 4E – 4F), indicating that *Xrp1* turns on *esg* expression in EEs. In support of this, we found that *Xrp1* and *esg* expression in EEs was moderately but significantly positively correlated in _“_AstC.EE_”_ and _“_CCHa2.EE_”_ (Fig. S4C – S4D).

Additionally, EEs can affect ISC behavior via ligand production and feedback signaling (Amcheslavsky et al., 2014; Biteau & Jasper, 2014). To investigate whether *Xrp1* O/E in EEs influenced ISC activities, we examined the number of *esg* mRNA^+^ progenitor cells and the number of mitotic cells, visualized by anti-Phospho-Histone H3 (pH3) staining. We found that they remained unchanged by *Xrp1* O/E in EEs (Fig. S4E – S4F). This result precluded the likelihood of a paracrine signaling that could affect the ISC state at the time point of examination. Collectively, these results suggest that *Xrp1* O/E in EEs specifically upregulates progenitor-related genes in EEs, thereby forcing EEs to dedifferentiate to progenitor cells.

### Forced *Xrp1* expression promotes EE plasticity in a progenitor state-dependent manner

Since radiation promotes EE plasticity via *Xrp1* expression, whose upregulation activates progenitor-specific gene expression, we returned to the lineage trace system to assess whether upregulating *Xrp1* alone was sufficient to promote EE plasticity. We focused on a subset of *NPF*-expressing EEs. *Xrp1* O/E in *NPF*-expressing EEs for 7 days increased the number of past EEs, which was further enhanced by *Xrp1* O/E for 14 days (Fig. 5A – 5B), proving sufficiency in the role of *Xrp1* for promoting EE plasticity. *Xrp1* O/E-driven EE plasticity was reproduced in *Tk*-expressing EEs (Fig. S5A – S5B). Notably, *Xrp1* O/E-induced EE fate conversion was only observed in the R2a region, whereas in the R2b region, despite their presence, EEs merely decreased in number without any fate conversion (Fig. S5C – S5D).

**Figure 5.**
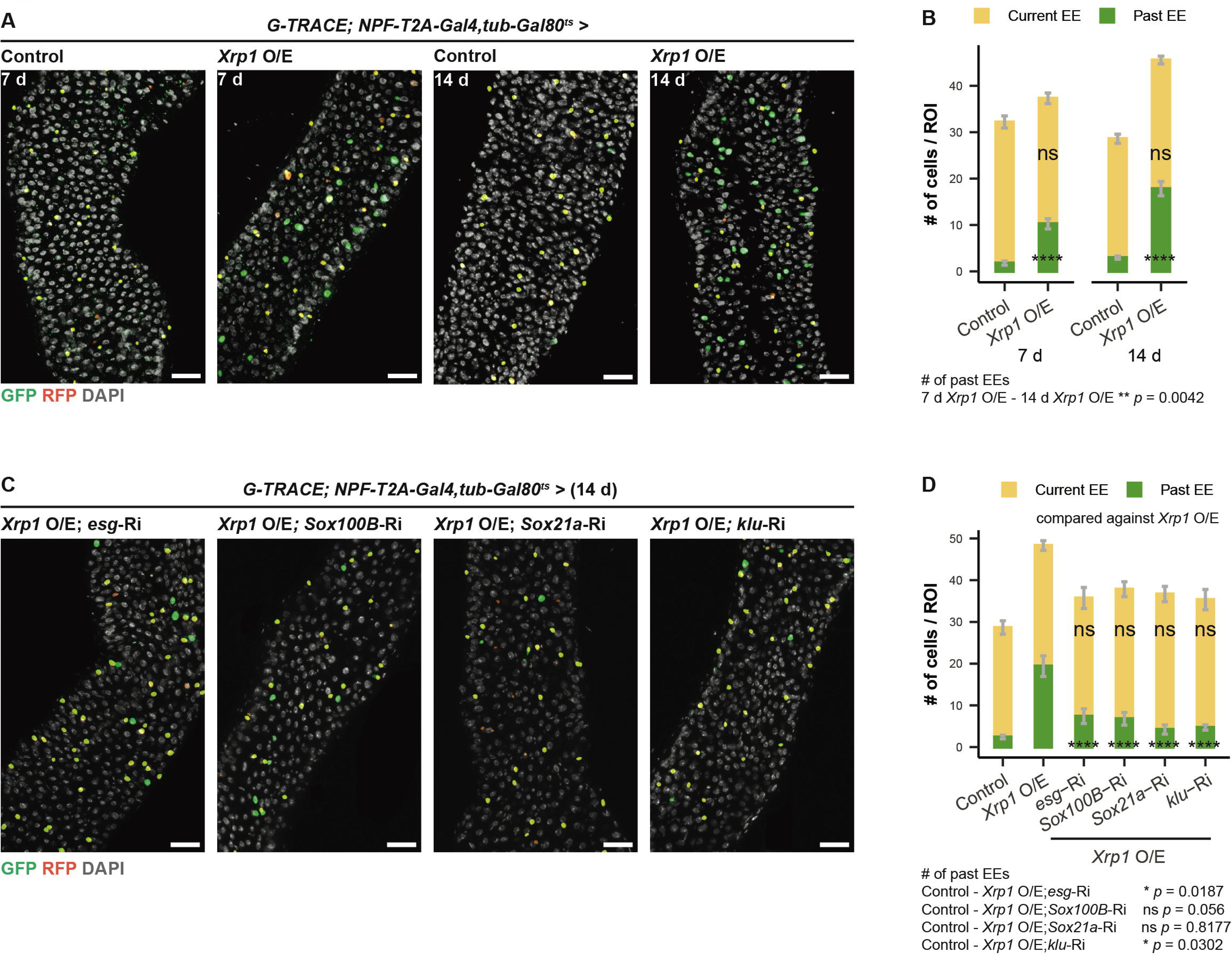
***Xrp1* O/E in EEs promotes EE plasticity in a progenitor state-dependent manner.** (A) Representative EE lineage-traced gut images of control and *Xrp*1 O/E in *NPF*-expressing EEs in 7-day-old and 14-day-old flies; stained with anti-GFP (green), anti-RFP (orange), and DAPI (white). Scale bars: 30 μm. (B) Quantification of the number of current EEs and past EEs corresponding to (A). N[gut] = 23 (7 d control), 30 (7 d *Xrp1* O/E), 56 (14 d control), 51 (14 d *Xrp1* O/E). Current EE: one-way ANOVA followed by Tukey HSD; Past EE: Kruskal-Wallis test followed by Steel-Dwass Test, **** *p* < 0.0001. (C) Representative EE lineage-traced gut images of *esg, Sox100B, Sox21a,* and *klu* knocked down on top of *Xrp1* O/E in *NPF*-expressing EEs in 14-day-old flies; stained with anti-GFP (green), anti-RFP (orange), and DAPI (white). Scale bars: 30 μm. (D) Quantification of the number of current EEs and past EEs, corresponding to (C). N[gut] = 21 (control), 22 (*Xrp1* O/E), 12 (*Xrp1* O/E; *esg-Ri*), 14 (*Xrp1* O/E; *Sox100B-Ri*), 13 (*Xrp1* O/E; *Sox21a-Ri*), 11 (*Xrp1* O/E; *klu-Ri*). Current EE: one-way ANOVA followed by Tukey HSD; Past EE: Kruskal-Wallis test followed by Steel-Dwass Test, **** *p* < 0.0001.

We also asked if there was any difference between the identity of past EEs derived from control EEs and *Xrp1* O/E EEs. Nuclear size measurement revealed two distinct peaks in past EEs from control guts, one coinciding with diploid EEs/ISCs and another approaching yet not reaching polyploid ECs. In comparison, the nuclear size of past EEs from *Xrp1* O/E EE showed a distribution shifting towards polyploid ECs, indicating a predominance of ECs (Fig. S5E). Consistent with their relatively large nucleus sizes, past EEs from *Xrp1* O/E EEs had a smaller chance of colocalizing with progenitor cell-expressed *esg* mRNA (Fig. S5F - S5G). In addition, some past EEs from *Xrp1* O/E EEs expressed the EB marker *NRE-LacZ* (Fig. S5H). These data suggest that *Xrp1* O/E alone can mimic the effect of radiation on EE plasticity, where EEs gained plasticity towards EC fate.

Having *Xrp1*-driven EE plasticity characterized, we asked whether the expression of progenitor-specific genes, such as *esg*, *Sox100B*, *Sox21a*, and *klu*, the potential downstream effectors (Fig. 4D, S4B, and S4C), were required for this process. Notably, we found that *Xrp1* O/E-driven EE plasticity was hindered when *esg*, *Sox100B*, *Sox21a,* and *klu* were knocked down, suggesting the necessity of going through a progenitor state in *Xrp1*-driven EE plasticity (Fig. 5C – 5D). Furthermore, EE-specific O/E of *Sox100B* and *Sox21a* increased the number of past EEs (Fig. S5I – S5J). These observations place *Xrp1* upstream of these cell fate determinants, which by themselves can affect EE plasticity as well. It can also be proposed that a portion of EEs, to an extent, have differentiation capacities and exhibit stem cell-like properties. Altogether, we present compelling evidence that *Xrp1* O/E in EEs promotes EE plasticity towards EC fate, in a manner dependent on progenitor cell fate determinants.

## Discussion

Here, we presented the first evidence of radiation-induced cellular plasticity of secretory EEs in the adult *Drosophila* midgut. Based on our findings, we proposed a model in which radiation triggers more EEs to dedifferentiate into ISCs, which then re-differentiate more preferably into ECs. We demonstrated that IR induced Xrp1 protein production in EE lineages; subsequently, *Xrp1* expression in EEs was found to be required for their radiation-induced plasticity. Mechanistically, *Xrp1* O/E in EEs led to ectopic expression of progenitor-specific gene *esg*, whose expression was necessary to confer the ability of *Xrp1* in promoting EE plasticity (Fig. 6). Our findings offer a mechanistic understanding of a form of cellular plasticity that represents a less well-characterized aspect of the epithelial damage response.

**Figure 6.**
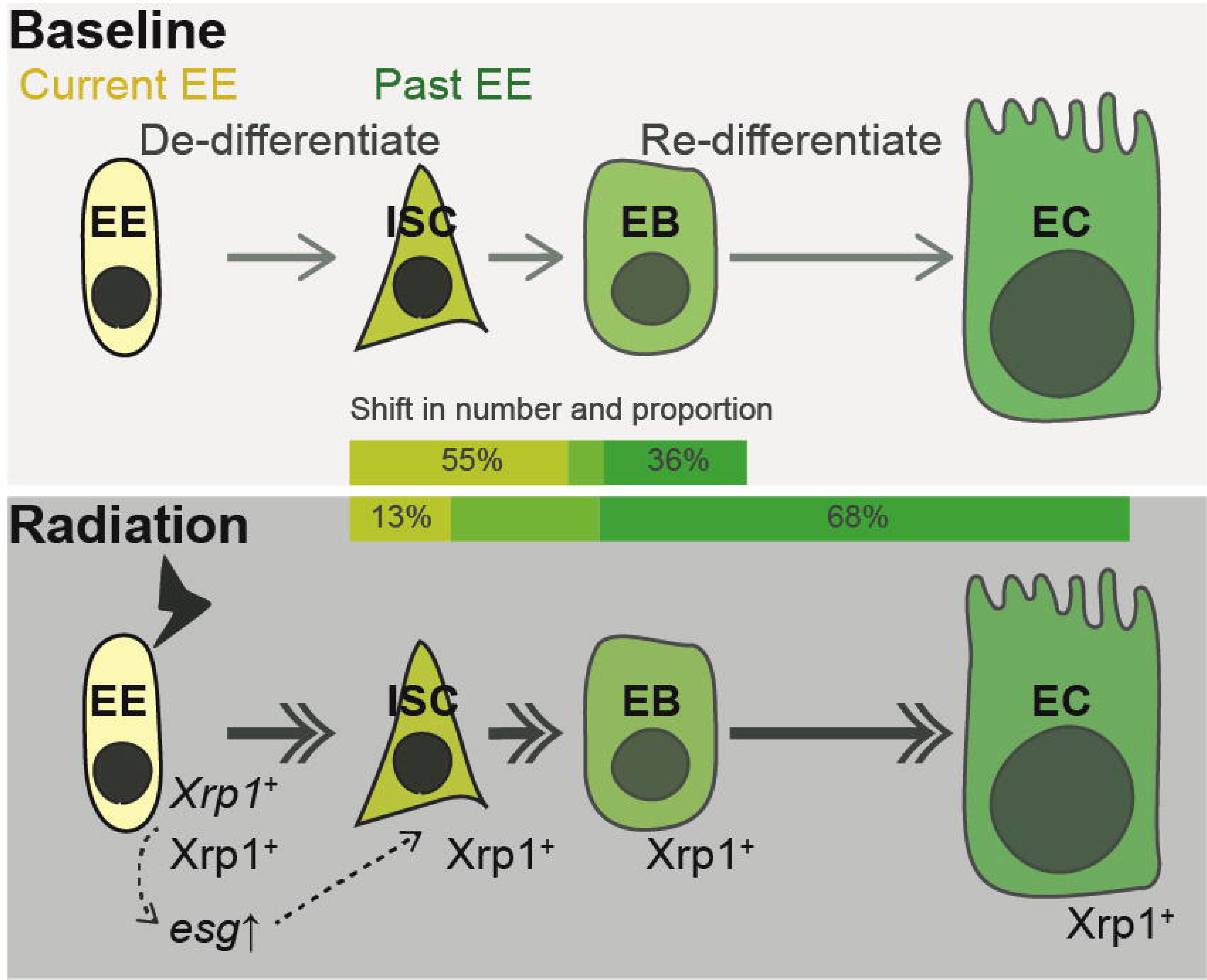
**Schematic of radiation-induced *Xrp1*-mediated EE plasticity.** While a small subset of EEs exhibits cellular plasticity under baseline conditions, radiation induces EEs to gain plasticity, which allows them to dedifferentiate into ISCs and re-differentiate into ECs. *Xrp1*, a stress-inducible transcription factor, is upregulated in EE lineages and is necessary for radiation-induced EE plasticity. Mechanistically, *Xrp1* induces ectopic expression of the progenitor-specific gene *esg*, whose expression is necessary for *Xrp1* to promote EE plasticity.

### *Xrp1*, a transcription factor implicated in cell differentiation

Extensive literature has reported *Xrp1* as a master regulator in maintaining genome stability upon DNA damage and eliminating competitively loser cells in the field of cell competition (Baillon et al., 2018; Blanco et al., 2020; Lee et al., 2016; Ochi et al., 2021). While *Xrp1* contributes to cell death in response to stress, we uncover a novel non-apoptotic role of *Xrp1* in intestinal cell differentiation. Combining findings from scRNA-seq and lineage tracing, our results demonstrate that *Xrp1* is sufficient to upregulate genes involved in the maintenance of progenitor identities and commitment to EC identities in EEs, thereby driving the identity transformation of EEs (Fig. 4D – 4F, S4C, and 5A – 5B).

Interestingly, the effect of *Xrp1* in promoting EE identity transformation is only observed in a specific region of the anterior midgut, while in another anterior midgut region, the death-promoting effect of *Xrp1* is more likely, given the decrease in the number of EEs that express *Xrp1* (Fig. S5C – S5D). The two distinct outcomes might be explained by EEs’ differences in the extent to which *Xrp1* can be overexpressed, or the basal expression level of certain Xrp1 targets or cofactors. Considering that the direct genomic targets of Xrp1 in wing imaginal discs and Xrp1-interacting proteins in in-vitro S2 cells have been reported (Baillon et al., 2018; Mallik et al., 2018), and EE transcriptomics have been resolved at a single-cell level, with spatial distribution of EE subtypes determined (Guo et al., 2019), it is worth exploring if EEs that can transform identity by *Xrp1* in the specific region have any differences in the expression of Xrp1-targerted genes. Additionally, the basal expression of *Xrp1* across all intestinal cell types might imply its contribution to other cellular processes beyond cellular plasticity. Although many details remain unclear, we propose that the transcription factor Xrp1 regulates progenitor-specific gene expression and, in combination with EE subtype–specific factors, drives the transformation of EE identity.

While the mammalian homolog of Xrp1 does not exist, Xrp1 is functionally homologous to C/EBP Homologous Protein, CHOP, in mammals (Baillon et al., 2018; Blanco et al., 2020). Similar as Xrp1, CHOP is a transcription factor induced as a result of integrated stress response or unfolded protein response, which promotes cell death, inhibits cell cycle and repairs DNA damage (Ferdoush et al., 2024; Liu et al., 2024; Ma et al., 2002; Wang et al., 1998). Although our findings do not focus on this conserved stress-regulated pro-apoptotic function, given the functional similarities between Xrp1 and CHOP, it is plausible that CHOP also participates in transcriptional regulation of cell differentiation. Importantly, CHOP is upregulated in liver tumors, where it promotes inflammation and fibrosis, thereby contributing to liver cancer (DeZwaan-McCabe et al., 2013). Our findings may thus suggest previously unrecognized aspects of pathogenetic pathways involving cell differentiation.

### *Xrp1*-mediated EE plasticity, an epithelial damage response program

Why differentiated cells bother to dedifferentiate has persisted as a debate over the years. A straightforward hypothesis to consider is that there is an imbalance between the proliferative capacity of stem cells and the growth demand of a tissue, which has been validated during regeneration after injury in the liver and intestine in mammals (Adkins-Threats & Mills, 2022; Li et al., 2020). In both mammals and fruit flies, following radiation damage, the loss of ISCs, the inability of ISCs to proliferate, and the increased gut permeability have been explicitly noted (Leibowitz et al., 2014; Pyo et al., 2014; Qiu et al., 2008; Sharma et al., 2020). Building upon these findings, it is conceivable that the plastic response that we observed (Fig. 2B – 2C) serves as another means to repopulate intestinal epithelial cells.

While we propose that radiation induces EEs to revert to progenitors, what confounds this hypothesis is the observation that EEs more favorably adopt an EC fate after radiation (Fig. 2D – 2E and S2D). Although we cannot rule out an alternative explanation for EE-EC trans-differentiation, our detailed time course analysis (Fig. 2D – 2E) leads us to assume that accelerated dedifferentiation to ISCs and re-differentiation to ECs may underlie the observed enhancement of EE plasticity.

In addition, the transient nature of EE identity transformation towards EC fate that we observed (Fig. 2C) is particularly interesting. A plausible hypothesis that aligns with the previously proposed model is that transiently present cells express mitogens that promote the proliferation and differentiation of surrounding progenitor cells, after which they undergo apoptosis, a phenomenon known as apoptosis-induced compensatory proliferation (Fogarty & Bergmann, 2017; Mollereau et al., 2013). This hypothesis is further supported by the presence of transiently undead or dying ECs that generate cytokine Unpaired 3 (Upd3) to promote ISC proliferation (Amcheslavsky et al., 2020; Jiang et al., 2011; Liang et al., 2017). Moving forward, it is essential to investigate whether the temporal appearance of ECs is needed for ISC activities or if the ISC pool maintenance affects EE plasticity.

Importantly, we uncover the requirement for *Xrp1* upregulation in EEs for their radiation-induced plasticity (Fig. 3D–3E). Although no clear homolog of *Xrp1* exists in vertebrates, p53, a widely conserved tumor suppressor gene, fulfills a similar role in mammals. A recent study has found that p53 is specifically enriched in revival stem cells in the mouse small intestine, which are thought to be dedifferentiated from progenies and are critically required for the emergence of this cell population after X-ray irradiation; without them, epithelial regeneration is impaired (Morral et al., 2024). Of particular note, p53 is required for *Xrp1* upregulation in *Drosophila* embryos and wing discs following X-ray irradiation (Akdemir et al., 2007; Baker et al., 2019; Brodsky et al., 2004; Khan & Baker, 2022). Our *Drosophila* study indicates that *Xrp1* mirrors how p53 reprograms irradiated intestinal epithelial cells in mammals, pointing to functional conservation at the level of biological process. Altogether, our work provides insights into the molecular underpinnings of damage-induced epithelial cellular plasticity.

## Materials and Methods

### Fly stocks and husbandry

Flies were maintained on a food made from agar (5.5 g/L), dry yeast extract (40 g/L), cornmeal (90 g/L), and glucose (100 g/L), heated until fully dissolved and mixed before adding propionic acid (3 mL/L) and 10 % butylparaben in 70 % ethanol (3.5 mL/L).

Flies were kept at 18 °C, 25 °C, or 29 °C in a 12 h:12 h light/dark cycle. Experimental crosses that did not involve Gal80^ts^-mediated manipulation were performed at 25 °C. When Gal80^ts^ was used, experimental crosses were raised at 18 °C. Newly eclosed virgin female adults were collected, maintained 0-2 days at 18 °C, and thereafter shifted to 29 °C for either 7 days or 14 days as indicated. The genotypes used were listed in the table below.

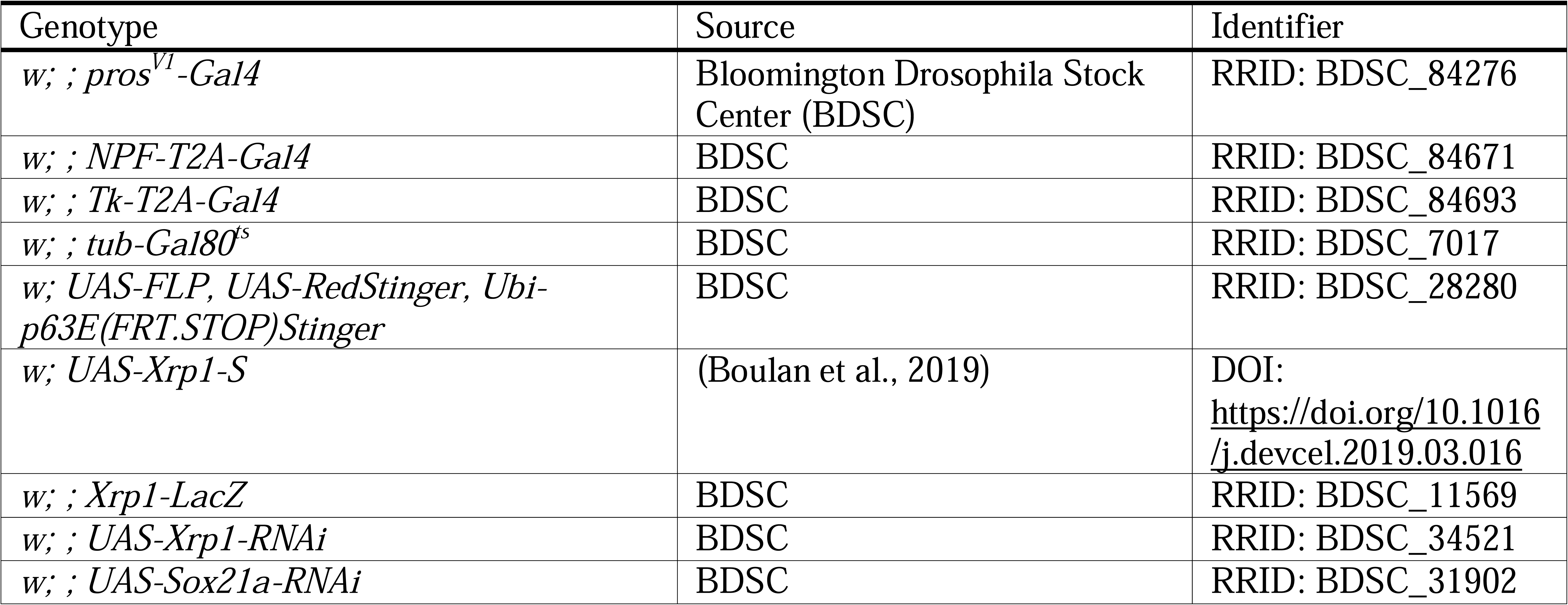
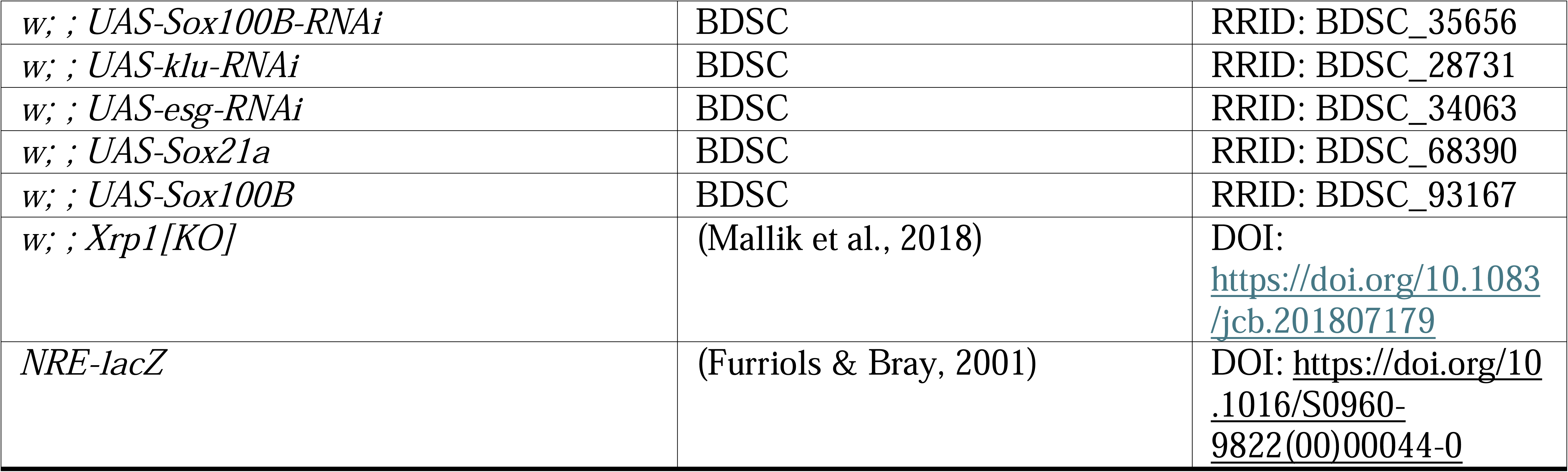

### Compound feeding

500 μl of different compound solutions were applied to vials containing 5 ml of 0.5 % agar at the bottom and tissues covering the surface of the agar. The vials were changed every 24 h. Flies were dry starved for 2 h on day 5 of EE lineage tracing, after which they were fed 5 % sucrose, or 5 % sucrose with 5 % DSS (Nacalai Tesque, Cat# 10912-92), or 10 mM Paraquat (Sigma-Aldrich, Cat# 856177) and sacrificed after 48 hours of feeding, on day 7 of EE lineage tracing.

### Ionizing radiation exposure

Adult flies were placed in vials with restricted space right before the experiment. Flies were exposed to 100 Gy X-ray irradiation, generated by the irradiator MX-160Labo, at 160 kV and 3 mA on day 6 of EE lineage tracing, and dissected several hours after X-ray irradiation as indicated in each figure. Non-irradiated control flies experienced the same amount of time in the irradiator that was turned off.

### Immunohistochemistry

Midguts were dissected in cold 1× PBS (Nacalai Tesque, Cat# 27575-31) and fixed in 4% paraformaldehyde (Nacalai Tesque, Cat# 02890-45) in PBS for 30 to 40 min at room temperature (RT). Fixed samples were washed three times in PBT [1× PBS supplemented with 0.1 % Triton X-100 (Nacalai Tesque, Cat# 35501-15)]. Samples were dehydrated in a series of EtOH gradient washes ranging from 10 % to 90 % and then kept in 90 % EtOH for 20Cmin. After rehydration, samples were blocked in blocking solution [PBT with 2 % BSA (Sigma-Aldrich, Cat# A9647-1006)] for 1 h at RT and then incubated with a primary antibody diluted to their proper concentration in the blocking solution at 4 °C overnight (refer to the table below).

The next day, the primary antibodies were washed off with three rinses and then incubated for 20 min in PBT. Samples were then incubated for 2 h at RT, in fluorophore (Alexa Fluor 488, 546, 555, or 633)-conjugated secondary antibodies at a 1:200 concentration in a blocking solution (refer to the table below). Samples were incubated three times in PBT, each for 20 min. During the last wash, DAPI (1:10000) was added to visualize the nucleus. Samples were finally mounted onto the slide glass using FluorSave Reagent (Merck Millipore, Cat# 345789).

Samples were visualized using a Zeiss LSM 900 confocal microscope. Z-stack images were captured at a 1024×1024 resolution. To count the cell numbers of current and past EEs, one image was taken per gut covering both the front and back epithelial layers using a 20× objective lens. For measuring the nucleus area and examining the colocalization, one or two images per gut covering the front epithelial layer were taken using a 40× objective lens with an AiryScan detector.

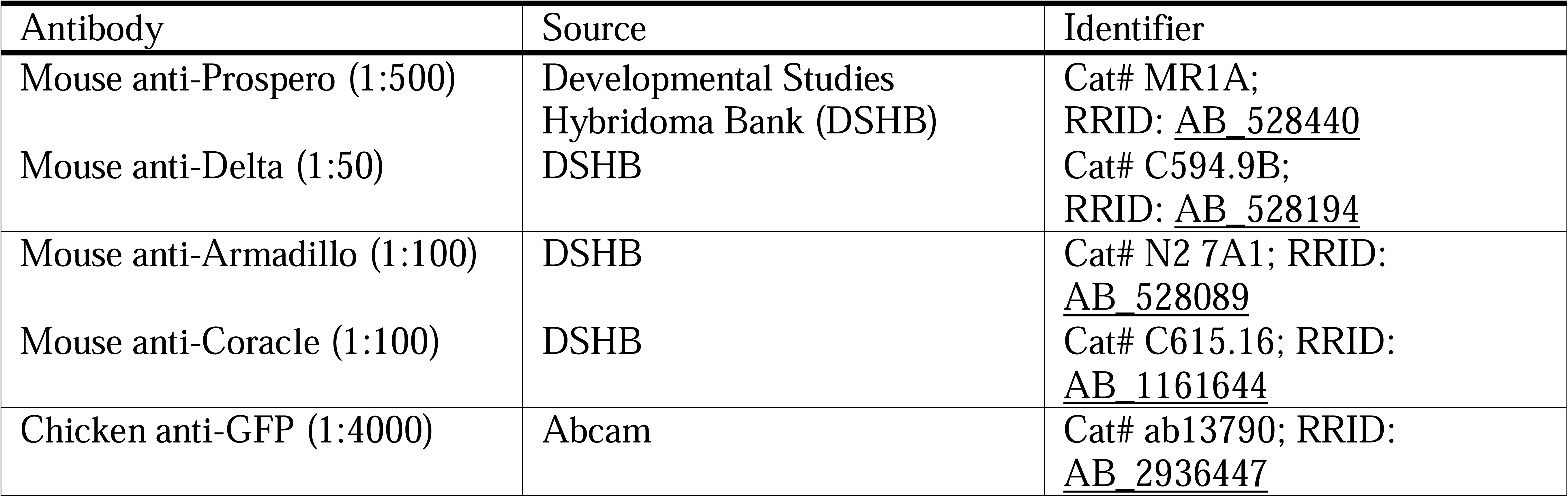
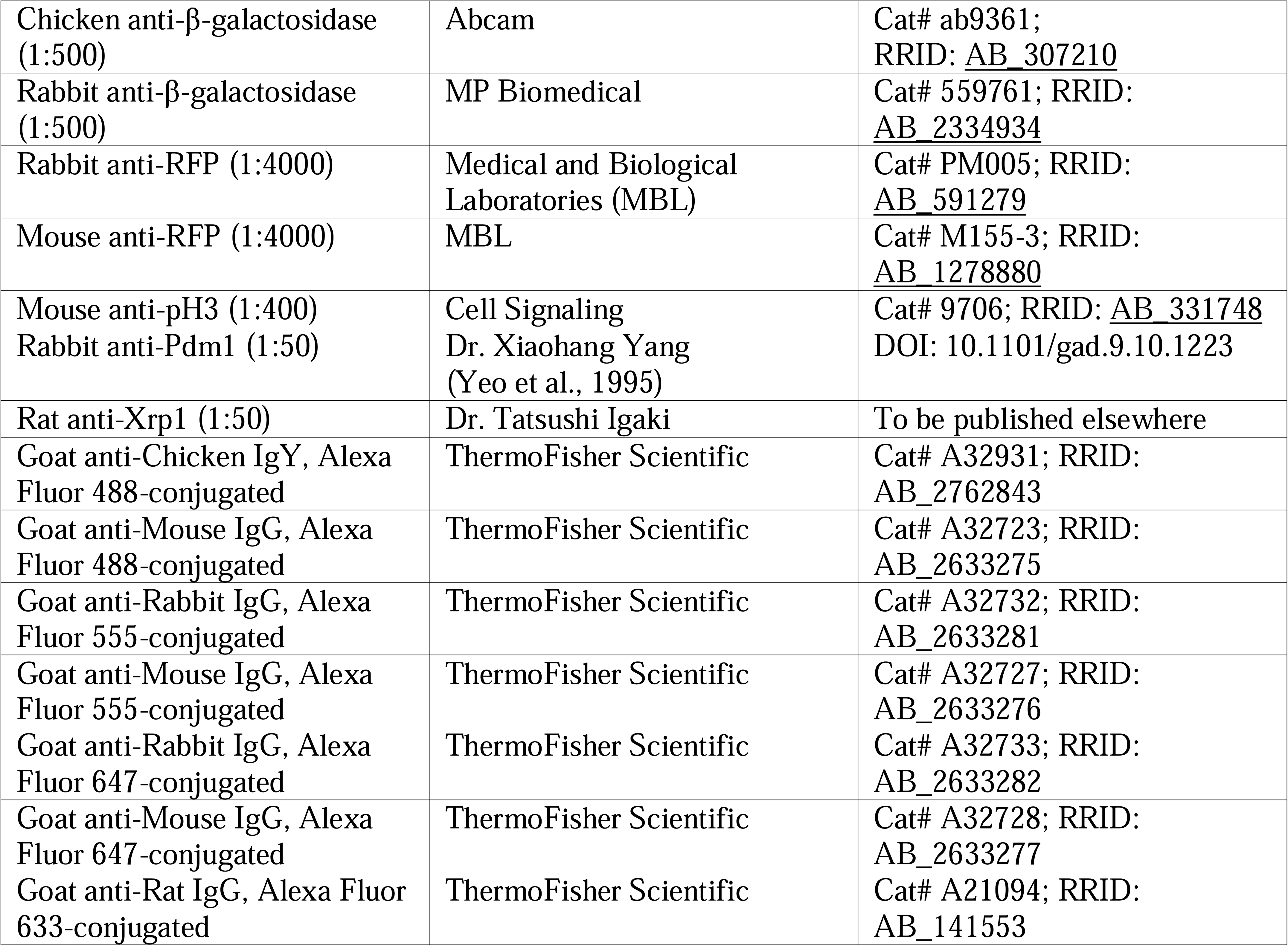

| Antibody | Source | Identifier |
| --- | --- | --- |
| Mouse anti-Prospero (1:500) | Developmental Studies Hybridoma Bank (DSHB) | Cat# MR1A; RRID: AB_528440 |
| Mouse anti-Delta (1:50) | DSHB | Cat# C594.9B; RRID: AB_528194 |
| Mouse anti-Armadillo (1:100) | DSHB | Cat# N2 7A1; RRID: AB_528089 |
| Mouse anti-Coracle (1:100) | DSHB | Cat# C615.16; RRID: AB_1161644 |
| Chicken anti-GFP (1:4000) | Abcam | Cat# ab13790; RRID: AB_2936447 |
| Chicken anti-β-galactosidase (1:500) | Abcam | Cat# ab9361; RRID: AB_307210 |
| Rabbit anti-β-galactosidase (1:500) | MP Biomedical | Cat# 559761; RRID: AB_2334934 |
| Rabbit anti-RFP (1:4000) | Medical and Biological Laboratories (MBL) | Cat# PM005; RRID: AB_591279 |
| Mouse anti-RFP (1:4000) | MBL | Cat# M155-3; RRID: AB_1278880 |
| Mouse anti-pH3 (1:400)<br>Rabbit anti-Pdm1 (1:50) | Cell Signaling<br>Dr. Xiaohang Yang<br>(Yeo et al., 1995) | Cat# 9706; RRID: AB_331748<br>DOI: 10.1101/gad.9.10.1223 |
| Rat anti-Xrp1 (1:50) | Dr. Tatsushi Igaki | To be published elsewhere |
| Goat anti-Chicken IgY, Alexa Fluor 488-conjugated | ThermoFisher Scientific | Cat# A32931; RRID: AB_2762843 |
| Goat anti-Mouse IgG, Alexa Fluor 488-conjugated | ThermoFisher Scientific | Cat# A32723; RRID: AB_2633275 |
| Goat anti-Rabbit IgG, Alexa Fluor 555-conjugated | ThermoFisher Scientific | Cat# A32732; RRID: AB_2633281 |
| Goat anti-Mouse IgG, Alexa Fluor 555-conjugated | ThermoFisher Scientific | Cat# A32727; RRID: AB_2633276 |
| Goat anti-Rabbit IgG, Alexa Fluor 647-conjugated | ThermoFisher Scientific | Cat# A32733; RRID: AB_2633282 |
| Goat anti-Mouse IgG, Alexa Fluor 647-conjugated | ThermoFisher Scientific | Cat# A32728; RRID: AB_2633277 |
| Goat anti-Rat IgG, Alexa Fluor 633-conjugated | ThermoFisher Scientific | Cat# A21094; RRID: AB_141553 |

### In situ hybridization chain reaction (isHCR)

We referred to a previous study (Mikami et al., 2025) for probe design and isHCR, with a few modifications. The probes used in this study are described below.

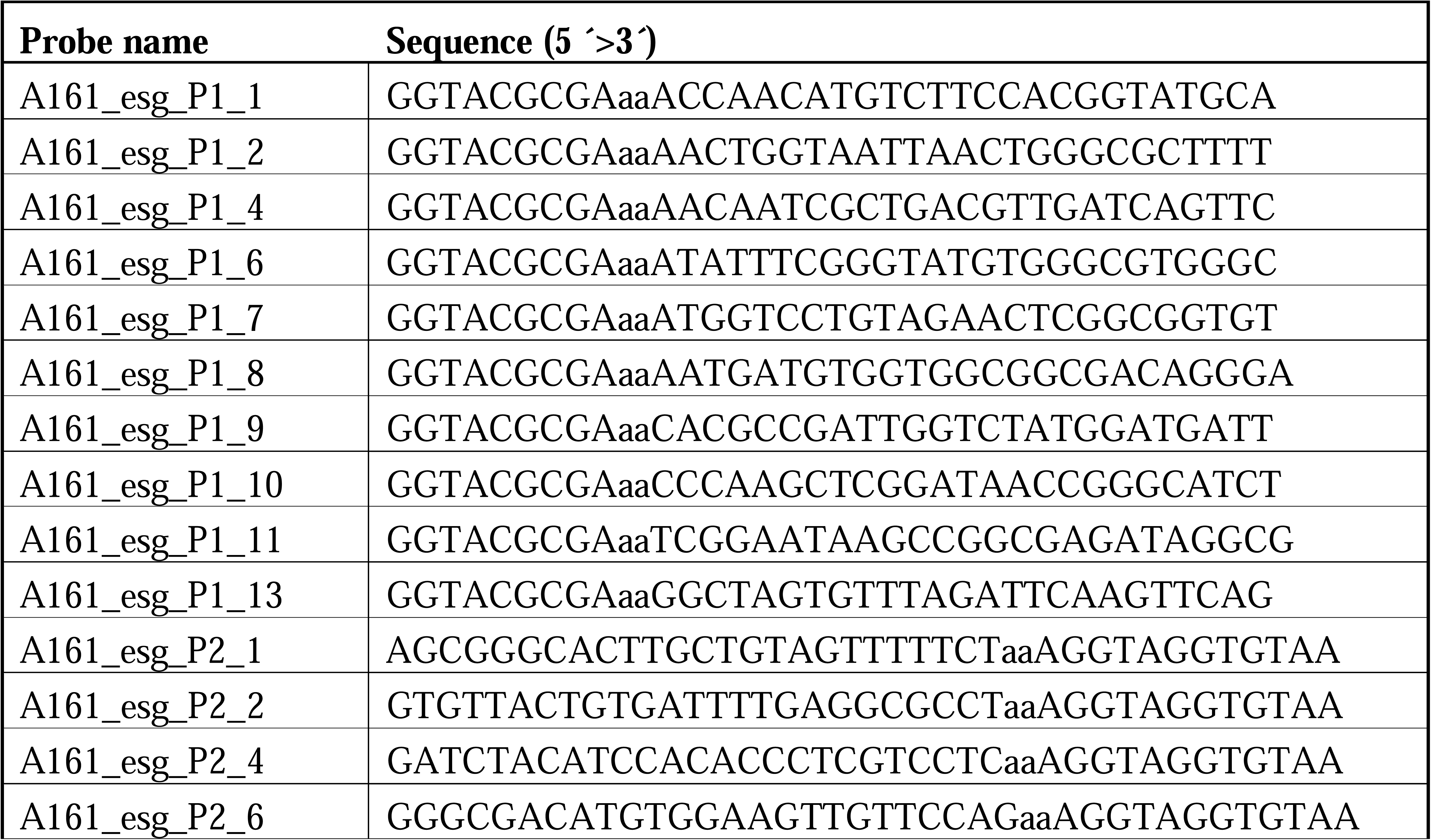
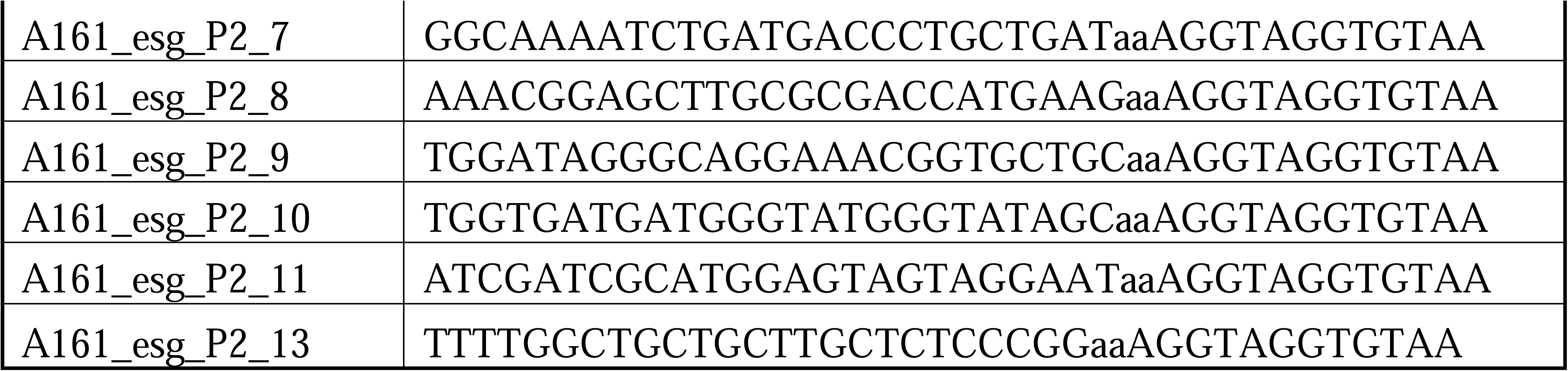

Dissected midguts were fixed with 4 % PFA for 30 min at RT, rinsed with PBT three times, sequentially dehydrated in 1:3, 1:1, and 3:1 PBT:MeOH, and stored in MeOH at - 20 °C. On the next day, midguts were sequentially rinsed with 3:1, 1:1, and 1:3 MeOH:1× SCCT [0.1 % Tween-20 (Fujifilm-Wako, Cat# 167-11515) in 1× SSC (150CmM Sodium Chloride (Nacalai Tesque, Cat# 31319-45) and 15 mM Trisodium Citrate Dihydrate, pH 7.0 (Fujifilm-Wako, Cat # 191-01785))], and washed with 1× SCCT for 3× 10 min. Midguts were pre-hybridized in pre-warmed hybridization buffer [15 % ethylene carbonate (Sigma-Aldrich, Cat# E26258), 5× SSC, 0.1 % Tween-20, 50 μg/ml heparin (Sigma-Aldrich, Cat# H3393-25KU), 1×Denhardt’s solution (Fujifilm-Wako, Cat# 043-21871)] for 30 min at 45°C. After this, midguts were hybridized in pre-warmed probe solution for 20 hours at 45°C. The probe solution was made by diluting each probe set to a final concentration of 20 nM in the hybridization buffer.

On the following day, midguts were washed with hybridization buffer for 2× 10 min at 45°C, rinsed with 1:1 hybridization buffer:1× SCCT, and lastly washed with 1× SCCT for 2× 10 min at RT. For the amplification reaction, A161 hairpin DNA pairs (H1 and H2, Nepagene, Cat# IPL-B-C-A161) were used. The hairpin DNA pairs were separately heated to 95 °C for 2Cmin and gradually cooled to 65 °C for 15 min and to 25 °C for 40 min before use. Midguts were incubated with amplification buffer (Nepagene, Cat# IPL-AB) with H1 and H2 hairpin DNAs at a 1:50 concentration, for 2 hours at 25°C. After this, midguts were rinsed with 1× SCCT, followed by 1:1 1× SCCT:PBT, and lastly with PBT three times. The midguts were then incubated in blocking solution for immunohistochemistry.

### Sample preparation for scRNA-seq

Bare midguts were dissected out in cold 1× Dulbecco’s phosphate-buffered saline (DPBS, ThermoFisher Scientific, Cat# 14200075) and transferred to a DNA Lo-bind tube containing DPBS on ice. After all the dissection was finished, the residual DPBS was removed from the tube, and 500 μl of 1mg/ml elastase (Sigma-Aldrich, Cat# E0258) in 1× DPBS was then added to the samples, which were incubated on a rotator at 27 °C. The samples were agitated by pipetting up and down ten times with wide-bore low-retention tips every 15 min. After 30 min, the supernatant was filtered through a 100 μm cell strainer into a new tube on ice, into which an equal amount of 1× DPBS with 1 % BSA was added to stop dissociation and prevent aggregation. Another 500 μl of 1mg/ml elastase in 1× DPBS was added to the undissociated guts for another round of dissociation. The cell suspension was pelleted at 200 x *g* for 5 min at 4°C. Cells were pooled together, resuspended in 300 μl of 1× DPBS with 0.1 % ultra-pure BSA (ThermoFisher Scientific, AM2616), and filtered through a 70 μm cell strainer.

For EE lineage-traced guts, GFP-positive cells were sorted out by FACS (MoFlo XDP, Beckman Coulter). Forward scatter and side scatter were used to sort out debris and doublets. Non-fluorescent wild-type flies were used to set negative fluorescence gates. Cells were collected into a DNA LoBind tube containing 1× DPBS with 0.1 % ultra-pure BSA.

Cell viability was assessed using an automated cell counter (Counteless, ThermoFisher) by mixing the cell suspension with trypan blue stain (ThermoFisher Scientific, Cat# 2524164). The cells (70 cells/μl, 60 % viability for FACS-sorted lineage-traced cells; 220 cells/μl, 83 % viability for lineage-traced all gut cells; 100 cells/μl, 78 % viability for control guts and 130 cells/μl, 84 % viability for *Xrp1* O/E EE guts) were then immediately applied to the 10X Genomics Chromium program using Chromium Next GEM Single Cell 3C Reagent Kits v3.1 (Cat# PN-1000128). Ten thousand cells were targeted for capture per sample and thus sequenced at a depth of 500 million paired-end reads with the Illumina Novaseq 6000.

### scRNA-seq analysis

The demultiplexing, barcoded processing, gene counting, and aggregation were made using the Cell Ranger software version 6.1.2. The Drosophila genome and annotation from the Berkeley Drosophila Genome Project, release 6 version 46 (BDGP6.46), were downloaded from the Ensembl Metazoa database (Yates et al., 2022). The sequences of marker genes, GFP (*pStinger*) and RFP (*RedStinger*), were obtained following the construction of G-TRACE (Evans et al., 2009), with *pStinger* originating from Stinger GFP transformation vector (GenBank: AF242364.2) and *RedStinger* originating from *Discosoma* drFP583 gene (GenBank: AF168419.2). They were added to FASTA and GTF to build a custom reference for Cell Ranger.

After obtaining the filtered barcodes, features, and matrix, we used the R package Seurat version 5.1.0 for analysis (Hao et al., 2024). Genes captured in less than 3 cells, and cells with less than 200 genes were removed. Cells with 500 ∼ 5000 genes detected and mitochondrial content smaller than 10 % were selected. Cells were normalized and rescaled by regressing out mitochondrial content using SCTransform version 0.4.1 (Hafemeister & Satija, 2019). A total of 2,564 cells from two datasets of FACS-enriched EE lineages and one dataset of all gut cells, and a total of 2,393 cells from control and *Xrp1* O/E EE guts were integrated using anchor-based integration. Dimensionality reduction was performed using RunPCA, and UMAPs were constructed using the top 30 principal components (McInnes et al., 2018). Clustering was performed using the Louvain algorithm at a resolution of 0.5.

Marker genes of each cluster were identified with a log fold change threshold of 0.5 and a minimum cell percentage of 25 %. Cell types were manually assigned to clusters based on the expression of characteristic marker genes (Guo et al., 2019; Hung et al., 2020): _“_ISC/EB_”_ named due to the progenitor marker *esg* expression, _“_EE_”_ named due to the EE marker *pros* expression, _“_EC_”_ due to the EC marker *nub* expression, _“_EC-like_”_ named due to its expression of the EC marker *nub* yet being far apart from other EC clusters by UMAP coordinates, _“_others_”_ named due to their similar gene expression profiles as the _“_ others_”_ in Hung’s dataset (Hung et al., 2020), and _“_ unk_”_ unknowable due to the absence of known marker expression. The scRNA-seq data reported in this study are available from Gene Expression Omnibus with accession code GSE301623.

### Quantitative and statistical analysis

Images were processed with Fiji/ImageJ (Schindelin et al., 2012). Maximum intensity projection was applied to each image. The cell number within each confocal image was manually counted. The nuclear area was measured by drawing around the border of the DAPI signal in the stacked 40× image.

All plots were generated with R (version 4.4.2). Bar plot data were shown as the mean ± standard error. Box plot data visualizes the median (thick line), first and third quartiles (box edges), and the minimum and maximum values (whiskers). Dots indicated individual values.

The Shapiro-Wilk test was applied to assess normality. When comparing two groups, an unpaired two-sided t test was used for parametric data, and the Wilcoxon Rank-Sum test was used for non-parametric data. When comparing multiple groups, one-way ANOVA was used for parametric data, and the Kruskal-Wallis test was used for non-parametric data. If the significance was obtained, post hoc tests were followed. For parametric data, Dunnett’s test (vs. control) or Tukey HSD (all pairs) was used; for non-parametric data, non-parametric

Dunnett’s test (vs. control) or Steel-Dwass test (all pairs) was used; * *p* < 0.05, ** *p* < 0.01, *** *p* < 0.001, **** *p* < 0.0001, not significant, (N.S.): *p* > 0.05.

## Raw data availability

All numerical raw data used for main figures and supplemental figures are described in Supplemental Data Files S1, S2, and S3.

## Acknowledgements

We thank Pierre Léopold, Tatsushi Igaki, Erik Storkebaum, Xiaohang Yang, the Bloomington Drosophila Stock Center, and the Developmental Studies Hybridoma Bank for providing us with fly stocks and other reagents. We are also grateful to Hiromi Yanagisawa, Satoru Kobayashi, Md Al Amin Sheikh, Maria Thea Rane Dela Cruz Clarin, and Yaxuan Cui for allowing us to use their equipment, and to Allison Bardin and Pierre Léopold for helpful discussions. This work was supported by grants from KAKENHI (JP22J01430 to H.N., 23K24025, 25H02543, 25K02406 to Y.N., and 22H00414 to R.N.), Japan Agency for Medical Research and Development AMED-CREST (20gm1110001 to R.N.), and the Cell Science Research Foundation (to Y.N.). Q.Q. was the recipient of the JST SPRING Grant Number JPMJSP2124.

## Author contributions

Conceptualization: Q.Q. and R.N. Methodology: Q.Q., H.N., Y.S., M.H., K.K., and R.N. Investigation: Q.Q. Visualization: Q.Q. Supervision: H.N., Y.S., Y.N., and R.N. Writing— original draft: Q.Q. Writing—review and editing: Q.Q., H.N., Y.S., M.H., K.K., Y.N., and R.N.

## Supplemental Figure Legends

**Figure S1. Detailed characterization of baseline EE plasticity, related to** Figure 1.

(A) Representative images of the whole EE lineage-traced midgut kept at restrictive temperature to prevent lineage tracing from being initiated; stained with anti-GFP (green), anti-RFP (orange), and DAPI (white). Scale bars: 100 μm.

(B) Top: Representative image of the whole EE lineage-traced midgut; stained with anti-GFP (green), anti-RFP (orange), and DAPI (white); gut regions indicated. Scale bar: 100 μm. Bottom: Representative EE lineage-traced gut images, from left to right, R2b, R3, and R4b; past EEs in R2b and R3 highlighted by arrows. Scale bars: 30 μm.

(C) Quantification of the proportion of past EEs (RFP^-^ GFP^+^ cells) among EE lineages (RFP^-^ GFP^+^, RFP^+^ GFP^-,^ and RFP^+^ GFP^+^ cells) per ROI. N[gut] = 32. Wilcoxon Rank-Sum test, *** *p* < 0.001.

(D, E, and F) Top: Representative gut image with *NPF*-expressing EEs lineage traced (D) and all EEs lineage-traced (E and F); all stained with anti-GFP (green), anti-RFP (orange), and DAPI (white); additionally, (D) anti-Pros (magenta; nucleus-localized) and anti-Dl (magenta; cytoplasm-localized), (E) anti-Arm (magenta), (F) anti-Cora (blue). Scale bars: 30 μm.

Bottom: Magnified views of cells indicated by numbered arrows on the image above. Top: merged; Bottom: (D) Pros and Dl; (E) Arm; (F) Cora. Scale bars: 3 μm.

(G) UMAP projection of individual cells from FACS-enriched EE lineages and unsorted all gut cells.

(H) Feature plot showing normalized expression of ISC/EB marker *esg*, *GFP*, EE marker *pros*, and *RFP* over UMAP projection of “ISC/EB”.

(I) Volcano plot showing genes differentially expressed between *GFP*^+^ and *GFP*^-^ ISC/EB, with each dot representing a gene, the x-axis showing log_2_ fold change, and the y-axis showing −log_10_ adjusted p-value. Raw data are available in Supplemental Data File S3.

**Figure S2. Detailed characterization of damage-induced EE plasticity, related to Figure 2**.

(A) Representative EE lineage-traced gut images of control, DSS, and paraquat-fed flies; stained with anti-GFP (green) and anti-RFP (orange). Scale bars: 30 μm.

(B) Quantification of the number of past EEs and current EEs, corresponding to (A). N[gut] = 20 (Control), 15 (DSS), 18 (Paraquat). Current EE: one-way ANOVA followed by Dunnett’s test, * *p* < 0.05, ** *p* < 0.01; past EE: Kruskal-Wallis test followed by Dunnett’s test.

(C) Quantification of the number of ISCs in non-irradiated (No IR) and irradiated (6 h, 10 h, 14 h, and 18 h Post IR) guts. N[image] = 28 (No IR), 24 (6 h Post IR), 24 (10 h Post IR), 34 (14 h Post IR), 24 (18 h Post IR). Kruskal-Wallis test followed by Dunnett’s test, * *p* < 0.05, ** *p* < 0.01, **** *p* < 0.0001.

(D) Quantification of the nucleus area of diploid EEs/ISCs, past EEs from non-irradiated (no IR) and irradiated (14 h Post IR) midguts, and polyploid ECs; illustrated by density plots showing the distribution and dashed lines indicating medians. n[cell] = 53 (No IR), 136 (14 h Post IR). n[cell] = 62 (EE/ISC), 67 (EC), measured in no-IR guts. Kruskal-Wallis test followed by Steel-Dwass Test.

**Figure S3. Detailed characterization of the effect of radiation associated with *Xrp1*, related to Figure 3**.

(A) Top: Representative gut images of non-irradiated (No IR) and irradiated (14 h Post IR) wild-type and *Xrp1* knock-out flies; stained with anti-Xrp1 (cyan) and DAPI (white). Bottom: Xrp1. Xrp1 protein highlighted by arrows. Scale bars: 30 μm.

(B) Top: Representative EE lineage-traced gut images of non-irradiated (No IR; top left) and irradiated (14 h Post IR; top right) flies; stained with anti-GFP (green), anti-RFP (orange), anti-Xrp1 (cyan), and DAPI (white). Scale bars: 30 μm.

Bottom: Magnified views of cells indicated by numbered arrows on the image above. Top: merged; Bottom: Xrp1. Scale bars: 3 μm.

(C) Top: Representative gut images of non-irradiated (No IR; top left) and irradiated (14 h Post IR; top right) flies with *Xrp1* transcript reported; stained with anti-β-Gal (green), anti-Pros (magenta; nucleus-localized), anti-Dl (magenta; cytoplasm-localized), and DAPI (white). Scale bars: 30 μm.

Bottom: Magnified views of cells indicated by numbered arrows on the image above. Top: merged; Bottom: β-Gal. Scale bars: 3 μm.

(D and E) Quantification of the ratio of β-Gal^+^ EEs among EEs per ROI (D) and the ratio of β-Gal^+^ ISCs among ISCs per ROI (E), corresponding to (C). N[gut] = 12 (No IR), 15 (14 h Post IR). Wilcoxon Rank-Sum test, **** *p* < 0.0001.

(F) Quantification of the ratio of Xrp1^+^ EEs among EEs per ROI. *EE > Control*: N[image] = 17 (No IR), 38 (14 h Post IR); *EE > Xrp1-Ri*: 12 (No IR), 28 (14 h Post IR). Kruskal-Wallis test followed by Steel-Dwass Test, ** *p* < 0.01. *EE > Control* No IR and 14 h Post IR data were reproduced from Fig. 3B to aid comparison.

(G) Representative EE lineage-traced gut images of non-irradiated (No IR; left) and irradiated (7 d Post IR; right) flies with and without *Xrp1* knocked down in EEs; stained with anti-GFP (green), anti-RFP (orange), and DAPI (white). Scale bars: 30 μm.

(H) Quantification of the number of past EEs and current EEs, corresponding to (G). *EE > Control*: N[gut] = 10 (No IR), 20 (7 d Post IR). *EE > Xrp1-Ri*: N[gut] = 16 (No IR), 28 (7 d Post IR). Current EE: One-way ANOVA test followed by Tukey HSD. ** *p* < 0.01.

**Figure S4. Detailed characterization of the effect of *Xrp1* O/E on EEs, related to Figure 4**.

(A) Violin plot showing normalized expression of EE-derived hormone peptides (*CCHa1*, *CCHa2*, *AstC*, *NPF,* and *Tk*) in each EE cluster split by the genotype.

(B) Violin plot showing normalized expression of cell type markers (*pros*, *esg*, *Dl*, *N*, and *klu*) in each EE cluster split by the genotype.

(C) Feature plot showing normalized expression of progenitor-related genes (*Sox100B*, *Sox21a,* and *klu*) in each EE cluster split by the genotype (left: control, right: *Xrp1* O/E EE; top: *Sox100B*, middle: *Sox21a*, bottom: *klu*).

(D) Feature plot showing normalized expression of *Xrp1* and *esg* in each EE cluster split by the genotype (left: control, right: *Xrp1* O/E EE; top: *Xrp1*, bottom: *esg*).

(E) Scatter plot with each dot representing each cell and dashed line indicating the linear regression model, where *Xrp1* counts and *esg* counts were positively correlated.

(F) Quantification of the number of *esg* mRNA^+^ cells per ROI. Wilcoxon Rank-Sum Test.

(G) Quantification of the number of mitotic (pH3^+^) cells per gut region. Wilcoxon Rank-Sum Test.

**Figure S5. Detailed characterization of the effect of *Xrp1* O/E on EE plasticity and the requirement of progenitor states, related to Figure 5**.

(A) Representative EE lineage-traced gut images of control and *Xrp*1 O/E in *Tk*-expressing EEs in 14-day-old flies; stained with anti-GFP (green), anti-RFP (orange), and DAPI (white). Scale bars: 30 μm.

(B) Quantification of the proportion of past EEs among EE lineages, corresponding to (A). N[gut] = 31 (control), 30 (*Xrp1* O/E). Current EE: unpaired two-sided t test; past EE: Wilcoxon Rank-Sum test, **** *p* < 0.0001.

(C) Representative EE lineage-traced gut images of control and *Xrp1* O/E in *NPF*-expressing EEs in the R2b region of 14-day-old flies; stained with anti-GFP (green), anti-RFP (orange), and DAPI (white). Scale bars: 30 μm.

(D) Quantification of the number of past EEs and current EEs, corresponding to (C). N[gut] = 47 (control), 42 (*Xrp1* O/E). Current EE: unpaired two-sided t test, **** *p* < 0.0001.

(E) Quantification of the nucleus area of diploid EEs/ISCs, past EEs from control and *Xrp1* O/E EE guts, and polyploid ECs in 7-day-old flies; illustrated by density plots showing the distribution and dashed lines indicating medians. n[cell] = 47 (control), 243 (*Xrp1* O/E). n[cell] = 52 (EE/ISC), 112 (EC), measured from control guts. Kruskal-Wallis test followed by Steel-Dwass Test.

(F and H) Top: Representative EE lineage-traced gut images of control (top left) and *Xrp1* O/E in *NPF*-expressing EEs (top right) in 7-day-old flies; all stained with anti-GFP (green), anti-RFP (orange) and DAPI (white), additionally, (D) *esg* mRNA (magenta), (E) anti-β-Gal (magenta). Scale bars: 30 μm.

Bottom: Magnified views of cells indicated by numbered arrows on the image above. Top: merged; Bottom: (F) *esg* mRNA; (G) β-Gal. Scale bars: 3 μm.

(G) The proportion of past EEs being small or large in nucleus size and being positive or negative for ISC/EB-expressed *esg* mRNA. Fisher’s exact test.

(I) Representative EE lineage-traced gut images of control and *Sox100B* and *Sox21a* O/E in *NPF*-expressing EEs in 14-day-old flies; stained with anti-GFP (green), anti-RFP (orange), and DAPI (white). Scale bars: 30 μm.

(J) Quantification of the number of current EEs and past EEs, corresponding to (I). N[gut] = 12 (control), 12 (*Sox100B* O/E), 18 (*Sox21a* O/E). Current EE: one-way ANOVA followed by Dunnett’s test; Past EE: Kruskal-Wallis test followed by Dunnett’s test, ** *p* < 0.01, *** *p* < 0.001.

## Supplemental Data Files (in Excel format)

**File S1. Raw data file for main figures**

**File S2. Raw data file for supplemental figures (except for Fig. S1I)**

**File S3. Raw data file for Supplemental Figure S1I.**

