## Supplementary figures and images for "*Xrp1* drives damage-induced cellular plasticity of enteroendocrine cells in the adult *Drosophila* midgut"

### Supplemental Figure S1

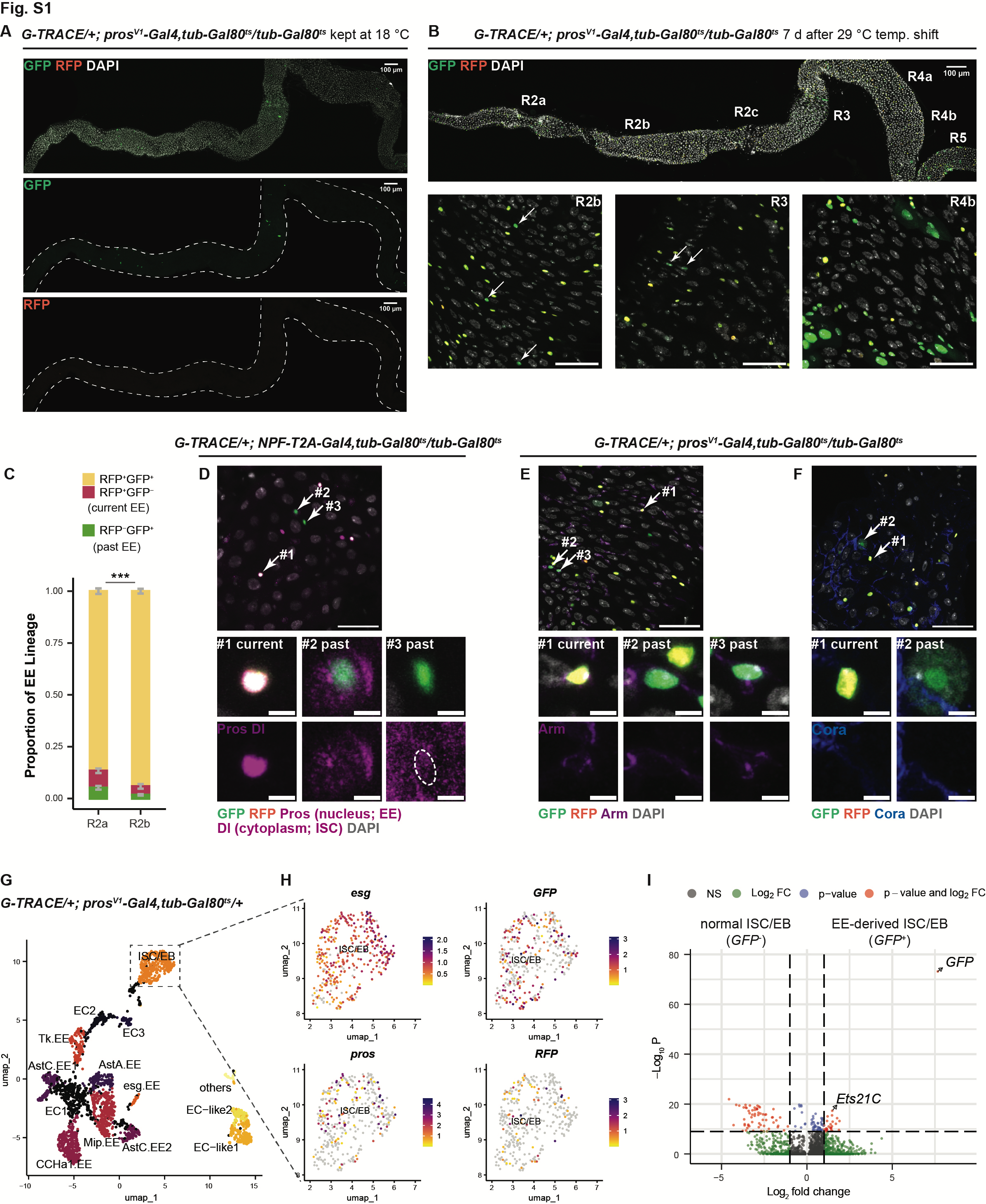

### Supplemental Figure S2

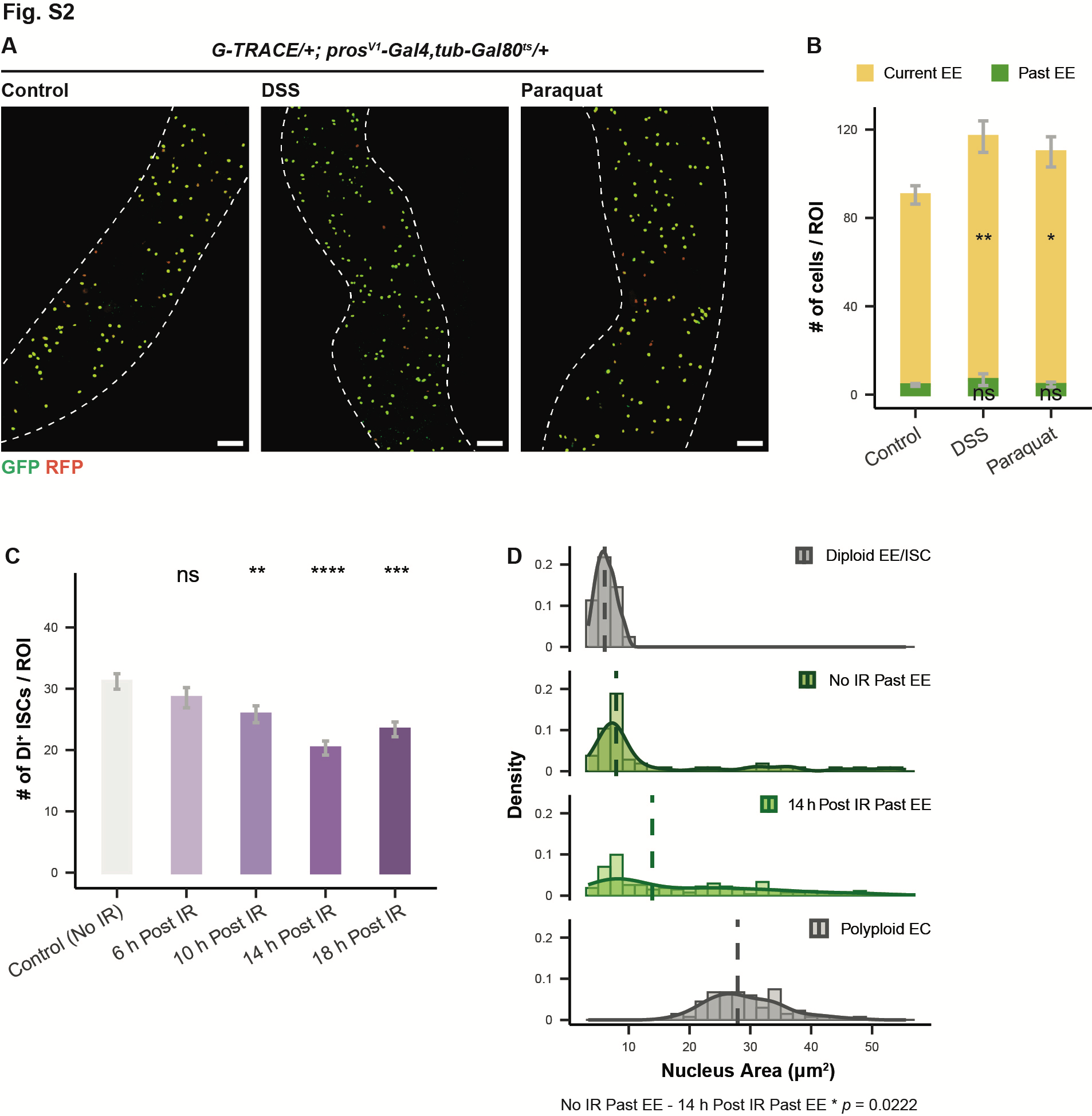

### Supplemental Figure S3

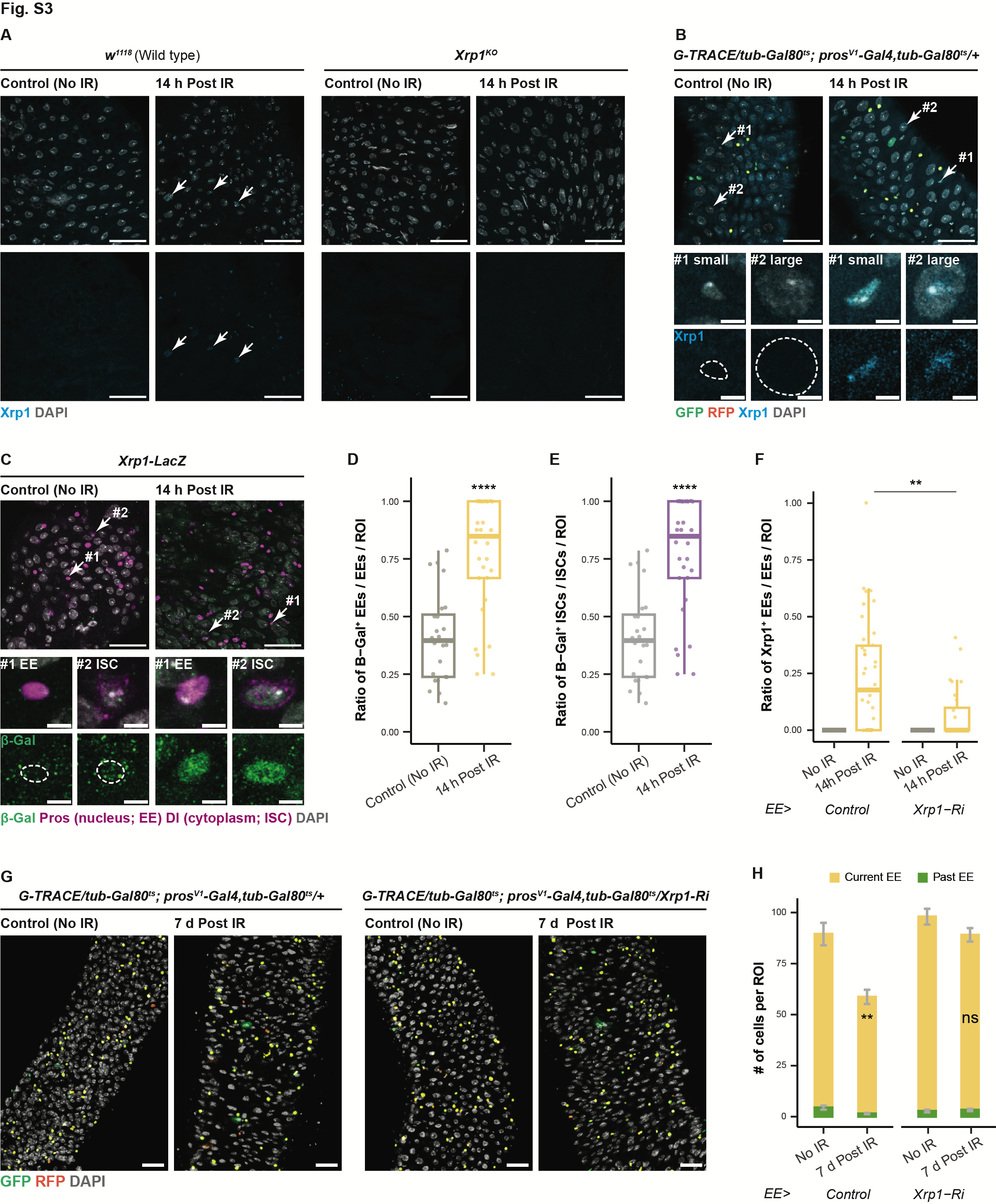

### Supplemental Figure S4

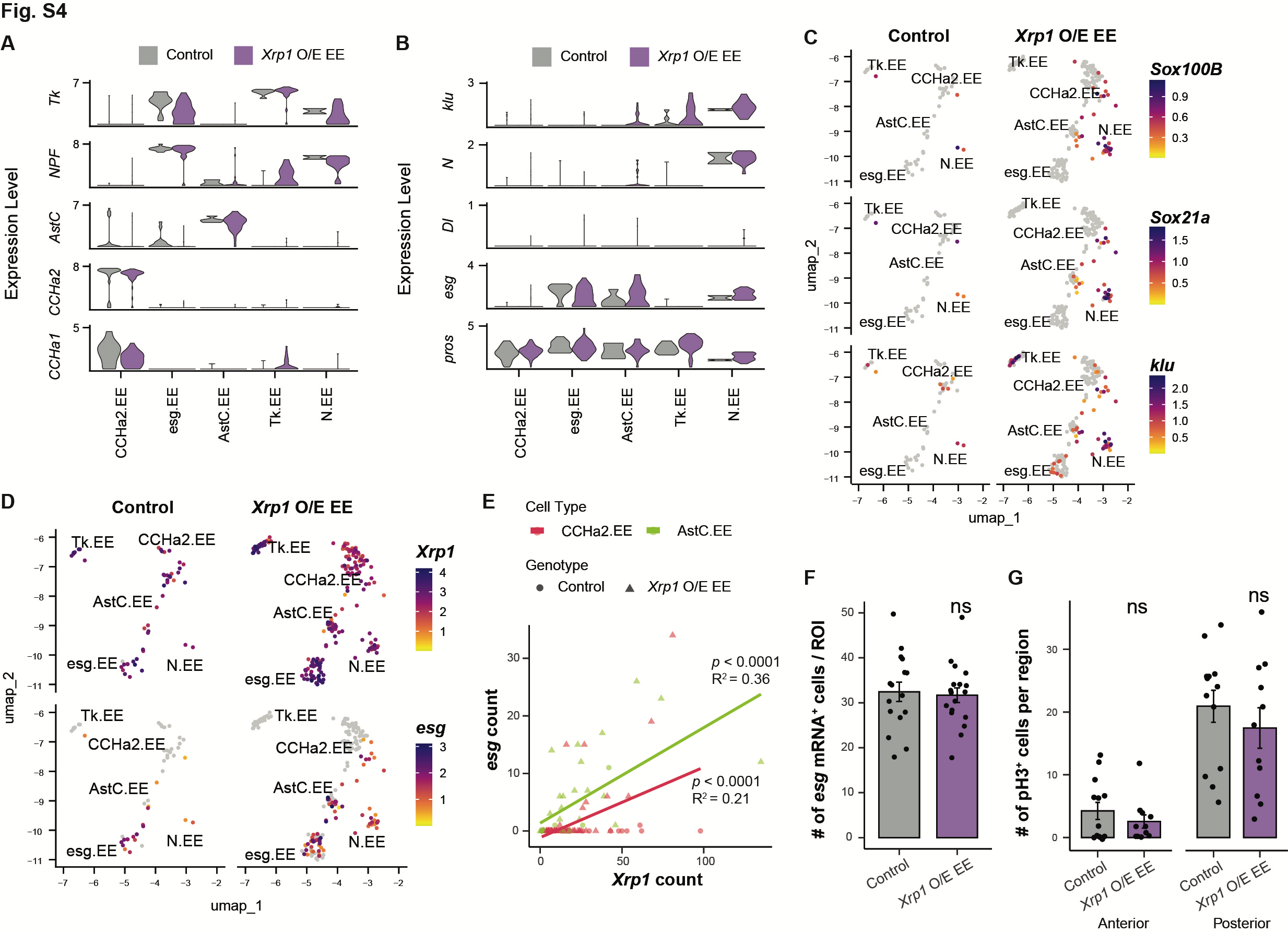

### Supplemental Figure S5

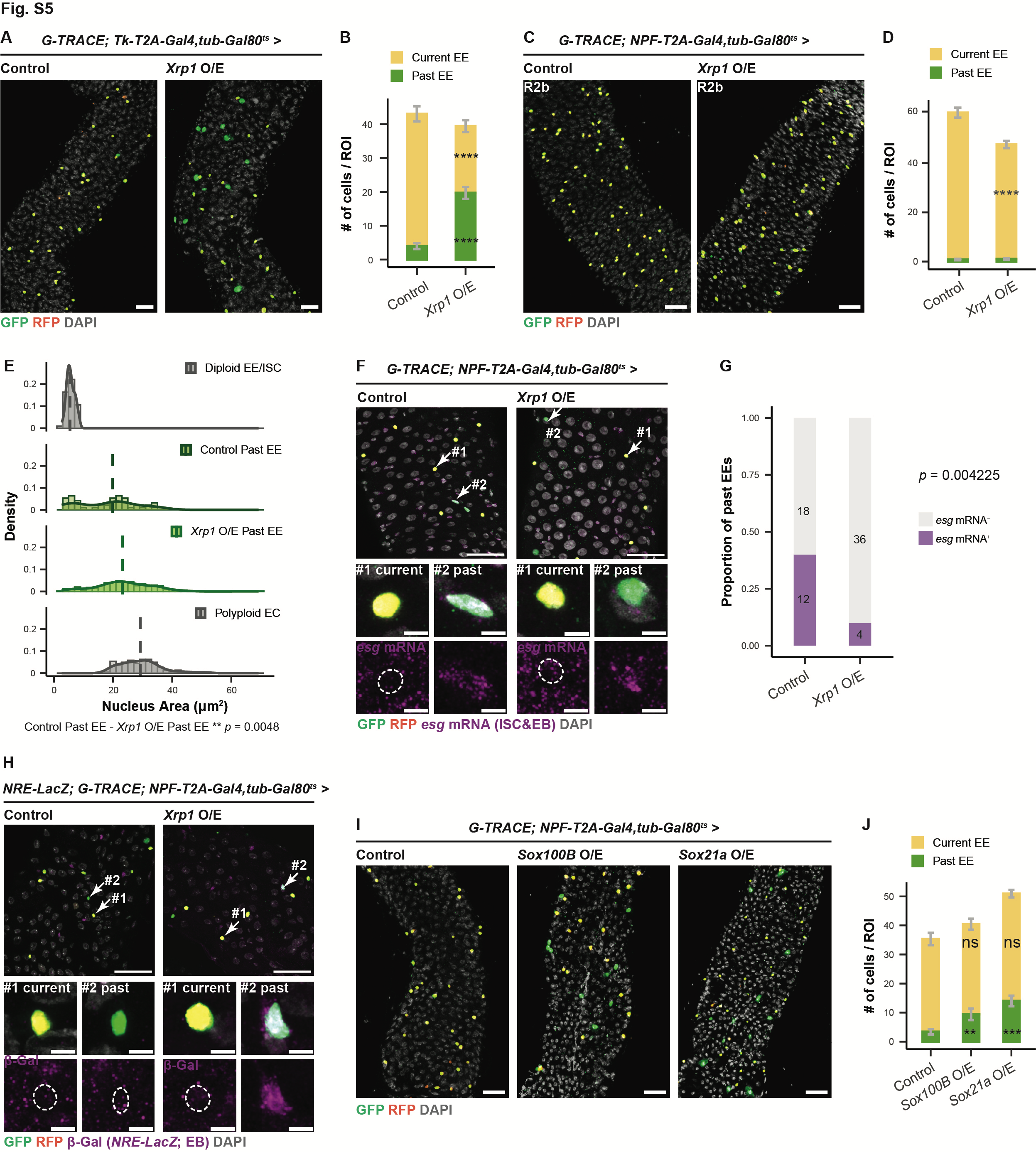
